# Mutations in the Bone Morphogenetic Protein signaling pathway sensitize zebrafish and humans to ethanol-induced jaw malformations

**DOI:** 10.1101/2023.06.28.546932

**Authors:** John R. Klem, Tae-Hwi Schwantes-An, Marco Abreu, Michael Suttie, Raeden Gray, Hieu Vo, Grace Conley, Tatiana M. Foroud, Leah Wetherill, CIFASD, C. Ben Lovely

## Abstract

Fetal Alcohol Spectrum Disorders (FASD) describe ethanol-induced developmental defects including craniofacial malformations. While ethanol-sensitive genetic mutations contribute to facial malformations, the impacted cellular mechanisms remain unknown. Bmp signaling is a key regulator of epithelial morphogenesis driving facial development, providing a possible ethanol-sensitive mechanism. We found that zebrafish mutants for Bmp signaling components are ethanol-sensitive and affect anterior pharyngeal endoderm shape and gene expression, indicating ethanol-induced malformations of the anterior pharyngeal endoderm cause facial malformations. Integrating FASD patient data, we provide the first evidence that variants in the human Bmp receptor gene *BMPR1B* associate with ethanol-related differences in jaw volume. Our results show that ethanol exposure disrupts proper morphogenesis of, and tissue interactions between, facial epithelia that mirror overall viscerocranial shape changes and are predictive for Bmp-ethanol associations in human jaw development. Our data provide a mechanistic paradigm linking ethanol to disrupted epithelial cell behaviors that underlie facial defects in FASD.

**Summary Statement:** In this study, we apply a unique combination of zebrafish-based approaches and human genetic and facial dysmorphology analyses to resolve the cellular mechanisms driven by the ethanol-sensitive Bmp pathway.

## Introduction

Ethanol is the most common environmental risk factor for congenital anomalies, with Fetal Alcohol Spectrum Disorders (FASD) describing all ethanol-induced birth defects. Global estimated incidence rates of FASD are .77%, though subpopulations of FASD can be much higher, ranging from 2-5% in the US and nearly 30% of individuals in some parts of the world with higher incidences of binge drinking (May et al., 2018; Popova et al., 2019). However, these numbers may be underestimates as nearly half of all pregnancies in the US are unplanned, and many pediatricians fail to recognize FASD (Finer & Zolna, 2016; Rojmahamongkol et al., 2015). Highly variable, multiple phenotypes present with FASD include structural malformations to the brain and face (Lovely, 2020). At the most severe end of the FASD spectrum is Fetal Alcohol Syndrome (FAS), which frequently presents as craniofacial defects including jaw hypoplasia (Blanck-Lubarsch et al., 2020). However, increasing data indicates that prenatal alcohol exposure (PAE) results in craniofacial defects in the absence of a diagnosis of FAS (Muggli et al., 2017; Suttie et al., 2013). Multiple factors contribute to the impact of PAE, in particular genetic predisposition (Lovely, 2020). To date, several ethanol-sensitizing alleles in zebrafish, mouse and humans have been linked with increased cell death, holoprosencephaly, oral clefting, disruption to axonal projections, and broad neural and eye defects (Boyles et al., 2010; Hong & Krauss, 2012; McCarthy et al., 2013; Swartz et al., 2014; Zhang et al., 2013). Despite these growing insights into the genetic components contributing to risk for FASD, we lack mechanistic insights into ethanol-sensitive gene function and associated cellular mechanisms during development that underlie FASD (Lovely, 2020).

Increasing evidence from several vertebrates including humans shows that genetic factors modulate developmental ethanol sensitivity (Lovely, 2020). Studying FASD in humans remains challenging due to the complex interplay of genetic background with often incompletely documented ethanol timing and dosage. Genetically tractable model organisms, such as zebrafish, have been essential in improving our understanding of the genetic loci behind the variability in FASD (Fernandes & Lovely, 2021). The zebrafish is well-suited for studying ethanol-sensitizing genetics because of its genetic tractability, high fecundity, external fertilization, embryo transparency, and rapid development (McCarthy et al., 2013; Swartz et al., 2014, 2020). We have previously used both a candidate-based and an un-biased forward-genetic screen-based approach to identify ethanol-sensitive mutations (McCarthy et al., 2013; Swartz et al., 2014, 2020) and these approaches have proven successful in predicting human gene-ethanol interactions (McCarthy et al., 2013). However, despite the increasing number of identified ethanol-sensitive loci, we lack a conceptual and mechanistic understanding of how these gene-ethanol interactions affect the diverse cell types and cellular behaviors underlying craniofacial development.

Previous work has established that both the genetic pathways required in, and the cellular events for, development of the craniofacial skeleton are deeply conserved between zebrafish and mammals (Knight & Schilling, 2006; Medeiros & Crump, 2012; Murillo-Rincón & Kaucka, 2020). Cranial neural crest cells (CNCC) give rise to the majority of the craniofacial skeleton and migrate from the dorsal neural tube to populate progenitor structures called the pharyngeal arches (Knight & Schilling, 2006; Medeiros & Crump, 2012; Murillo-Rincón & Kaucka, 2020). Concurrent with cranial neural crest cell migration, the pharyngeal endoderm undergoes its own cellular rearrangements and tissue movements to form a midline epithelial sheet with lateral protrusions called “pouches.” Proper morphogenesis of the pharyngeal endoderm is critical for craniofacial development, in particular of the jaw (Balczerski et al., 2012; Couly et al., 2002; Crump, 2004; Haworth et al., 2004, 2007; Lovely et al., 2016). Work in zebrafish has shown that several genetic pathways regulate endodermal morphogenesis (Balczerski et al., 2012; Choe et al., 2013; Choe & Crump, 2014; Crump, 2004; Hu et al., 2018; Li et al., 2019; Lovely et al., 2016). One such pathway is the Bone Morphogenetic Protein (Bmp) signaling pathway. Comprised of over 60 pathway components with different spatio-temporal expression and activity, active Bmp signaling initiates with heterodimer ligands binding to a complex of two type I and two type II transmembrane receptors that regulate downstream target genes through phosphorylation of Smad proteins (Kondo, 2007; Little & Mullins, 2009). Two major interacting ligands of the pathway are Bmp2b and Bmp4, both of which have a higher binding affinity type I Bmp receptors, such as Bmpr1bb over type II receptors (Little & Mullins, 2009; Tajer et al., 2021). We have previously shown that Bmp signaling is required in the endoderm to regulate a Fibroblast Growth Factor (Fgf) signaling response in the forming pouches (Lovely et al., 2016). Chemical inhibition of BMP signaling impairs the morphogenesis of both pouches and the anterior pharyngeal endoderm as the area of endoderm anterior to the first pouch, resulting in craniofacial malformations (Lovely et al., 2016). These observations showed that functional Bmp signaling is indispensable for establishing proper endoderm morphology as necessary for uninterrupted craniofacial skeleton patterning.

As a complex pathway that is essential for craniofacial morphogenesis, we hypothesized that BMP signaling is potentially ethanol sensitive. Here, we tested the ethanol sensitivity mutations in several components of the Bmp pathway in zebrafish. We show that hemi- or homozygous mutants for the Bmp ligand genes *bmp2b* and *bmp4* and for the receptor gene *bmpr1bb* (defined as Bmp mutants henceforth) predispose zebrafish embryos to distinct ethanol-induced craniofacial shape changes, particularly jaw malformations. Using quantitative morphometrics, we show that ethanol-induced disruptions to anterior pharyngeal endoderm shape, which mirror changes in facial shapes, alter the expression domain of the oral ectoderm marker *fgf8a* as associated with jaw malformations. We go on to show using fluorescent analyses that Bmp signaling response are lost specifically in the endoderm in Bmp mutants, but ethanol does not impact Bmp signaling in any meaningful way. Genetic analysis shows that *BMPR1B* associates with jaw deformations in children with ethanol exposure that mirror our zebrafish data, underlining the predictive strength of our zebrafish findings. Collectively, our data links perturbations in Bmp signaling to ethanol susceptibility during craniofacial development in zebrafish and humans, establishing mechanistic concepts in gene-ethanol interactions for future studies in FASD.

## Results

### Mutations in multiple components of the Bmp pathway sensitize embryos to ethanol-induced viscerocranial malformations

To build on previous work where we identified multiple ethanol sensitive genetic loci that predisposed to ethanol-induced craniofacial malformations (McCarthy et al., 2013; Swartz et al., 2014, 2020), we performed a candidate screen to identify additional ethanol-sensitive genes. From this screen, we identified the Bmp signaling pathway components, pathway ligands *bmp2b* and *bmp4* (Mullins et al., 1996; Stickney et al., 2007) and the pathway receptor *bmpr1bb* (Neumann et al., 2011) as being ethanol sensitive genes regulating facial development. Our previous work showed that Bmp signaling is required for facial development by regulating pharyngeal endoderm morphogenesis from 10-18 hours post fertilization (hpf) (Lovely et al., 2016). We also showed that the Bmp ligand genes *bmp2b* and *bmp4* are expressed adjacent to the development endoderm during this time window (Lovely et al., 2016). In addition, both of Bmp2b and Bmp4 have a higher binding affinity type I Bmp receptors over type II (Little & Mullins, 2009; Tajer et al., 2021), linking the *bmpr1bb*. This ultimately provides a testable mechanism for ethanol-induced craniofacial defects that we examine below.

To test these Bmp signaling mutants, we exposed embryos to a sub-phenotypic dose of 1% ethanol (v/v) originally from 6 hpf to 5 days post fertilization (dpf). We selected 1% ethanol (v/v) as the highest dose that does not cause craniofacial defects in wild type embryos, while higher doses of 1.25% and up have been shown to impact facial development (Bilotta et al., 2004; Everson et al., 2022; McCarthy et al., 2013; Swartz et al., 2014; Zhang et al., 2014). 1% ethanol (v/v) equilibrates within 5 minutes of exposure to an average ethanol tissue concentration of 50 mM (approximately 30% of the media) (Flentke et al., 2014; Lovely et al., 2014; Reimers et al., 2004; Zhang et al., 2013). Importantly, this embryonic ethanol concentration is roughly equivalent to a human Blood Alcohol Concentration of 0.23; while a binge dose, is physiologically relevant to FASD and humans are readily capable of surpassing this amount (Canfield et al., 2019; Ethen et al., 2009; Jones, 2008; Maier, 2001; Whaley et al., 2019).

Using this long exposure paradigm, we found that mutations in our chosen Bmp mutants sensitize developing zebrafish to a range of ethanol-induced developmental defects, including small eyes, defects to bone mineralization and, for this work, jaw malformations (Fig. 1A-H). While *bmp2b*^-/-^ embryos do not develop past 16 hpf (Nguyen et al., 1998), heterozygous *bmp2b^+/-^*, *bmp4^-/-^*, or *bmpr1bb^-/-^* larvae undergo normal craniofacial development and are superficially indistinguishable from their wild type siblings (Fig. 1A-D). However, when exposed to ethanol using our conditions of 1% ethanol (v/v), these Bmp mutants developed a range of jaw defects from malformations to outright absence of the jaw (Fig. 1 E-F). The expressivity of the spectrum of jaw phenotypes was consistent between the different mutant lines with only the penetrance changing between the lines (Fig. 1 E-F, Table 1). Jaw malformations were more common in ethanol-treated Bmp mutants than absent jaw, with 15.9% vs 4.6% in *bmp2b^+/-^*, 37.8% vs 6.13% in *bmp4^-/-^*, and 29.1% vs 2.7% in *bmpr1bb^-/-^*, with variation between experimental groups (Table 1). Our quantifications also demonstrated that while wild type siblings never displayed sensitivity to ethanol, heterozygous *bmp4^+/-^* and *bmpr1bb^+/-^* larva were ethanol sensitive with incomplete penetrance (Table 1). These results reveal that consistent viscerocranial malformations, in particular jaw absence, occur in ethanol-treated zebrafish with Bmp pathway mutations but not in either Bmp mutant or ethanol treatment alone.

**Figure 1.**
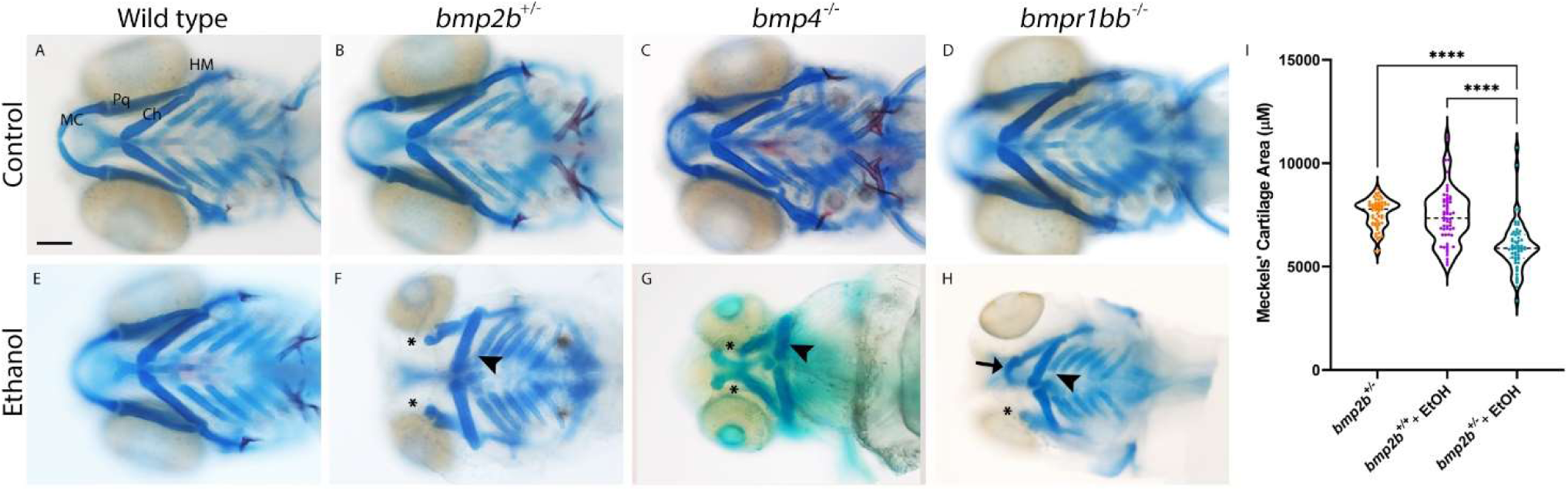
Multiple members of the Bmp pathway display ethanol sensitive facial phenotypes. (**A-H**) Whole-mount images of viscerocranium at 5 dpf larva. Cartilage is blue and bone is red (Ventral views, anterior to the left, scale bar: 100 μm, MC = Meckel’s, Pq = palatoquadrate, Ch = ceratohyal and HM = hyomandibular cartilages). (**I**) Area measures of the Meckel’s cartilage. (**A-D**) *bmp2b^+/-^* or *bmp4^-/-^* or *bmpr1bb^-/-^* larva develop comparable to Wild type larva. (**E-H**) Exposure of 1% ethanol exposure from 10-18 hpf results in a range of defects to the viscerocranium, from loss of MC at the extreme end of this range (asterisks) to reductions in size and changes in shape in the MC (arrows) as well as a flattening of the Ch (arrowheads), the average ethanol-induced defects, in the Bmp mutant alleles but not in Wild type siblings. (**I**) The size of Meckel’s cartilage elements was reduced in ethanol-treated *bmp2b^+/-^*larva compared to ethanol-treated wild type and untreated *bmp2b^+/-^*larva (F=36.85, p=.0001, one-way ANOVA, n=29 larvae, both Meckel’s cartilage elements per group (58 total)).

**Table 1.** Penetrance of gene-ethanol interactions in Bmp mutants. Percent of jaw malformations and jaw loss per experiment in ethanol-treated Bmp single mutant embryos generated from random heterozygous crosses.

| Experiments | Untreated Embryos | Jaw Malformations | Percent | EtOH-treated Embryos | Jaw Malformations | Percent | Jaw Loss | Percent |
| --- | --- | --- | --- | --- | --- | --- | --- | --- |
| *bmp2b* |  |  |  |  |  |  |  |  |
| 1 | 37 | 0 | 0.0% | 41 | 6 | 14.6% | 0 | 0.0% |
| 2 | 45 | 0 | 0.0% | 41 | 10 | 24.4% | 3 | 7.3% |
| 3 | 37 | 0 | 0.0% | 115 | 20 | 17.4% | 4 | 3.5% |
| 4 | 80 | 0 | 0.0% | 46 | 9 | 19.6% | 5 | 10.9% |
| 5 | 64 | 0 | 0.0% | 61 | 12 | 19.7% | 1 | 1.6% |
| 6 | 75 | 0 | 0.0% | 193 | 26 | 13.5% | 7 | 3.6% |
| 7 | 23 | 0 | 0.0% | 24 | 0 | 0.0% | 4 | 16.7% |
| Totals | 361 | 0 | 0.0% | 521 | 83 | 15.9% | 24 | 4.6% |
| *bmp4* |  |  |  |  |  |  |  |  |
| 1 | 24 | 0 | 0.0% | 63 | 30 | 47.6% | 3 | 4.8% |
| 2 | 41 | 0 | 0.0% | 131 | 38 | 29.0% | 2 | 1.5% |
| 3 | 42 | 0 | 0.0% | 38 | 24 | 63.2% | 2 | 5.3% |
| 4 | 51 | 0 | 0.0% | 94 | 20 | 21.3% | 16 | 17.0% |
| 5 | 44 | 0 | 0.0% | 82 | 42 | 51.2% | 2 | 2.4% |
| Totals | 202 | 0 | 0.0% | 408 | 154 | 37.7% | 25 | 6.1% |
| *bmpr1bb* |  |  |  |  |  |  |  |  |
| 1 | 105 | 0 | 0.0% | 82 | 41 | 50.0% | 1 | 1.2% |
| 2 | 48 | 0 | 0.0% | 38 | 1 | 2.6% | 0 | 0.0% |
| 3 | 154 | 0 | 0.0% | 69 | 13 | 18.8% | 4 | 5.8% |
| Totals | 307 | 0 | 0.0% | 189 | 55 | 29.1% | 5 | 2.6% |

From this initial long exposure paradigm, we narrowed down the exposure window from 10-18 hpf, (the same window when Bmp signaling is required for endoderm morphogenesis and jaw development (Lovely et al., 2016)), with exposure before 10 hpf not adding to the penetrance and expressivity of the phenotypic spectrum each single mutant (data not shown). We chose to work on *bmp4* larvae in all subsequent experiments as the mutation of *bmp2b* is weakly dominant (Kishimoto et al., 1997) and *bmpr1bb* is in the WIK background, whereas all of the other Bmp mutants used in this study are in the AB background, both of which our later analyses. In addition, we analyzed *bmp4* larvae as they showed the greatest variation due to ethanol treatment. (Table 1). To test the onset of ethanol-induced facial malformations, we started the ethanol exposure paradigm on wild type and *bmp4^-/-^* embryos at 10 hpf, 14 hpf and 18 hpf. Our analysis showed that as the onset of ethanol exposure started at later developmental staging the percent of jaw loss and jaw malformation decreased (Table S1). Starting our ethanol exposure paradigm at 24 hpf, we observed no jaw loss and only 1.2% of embryos with jaw malformations (Table S2). To test if dosage was the determining factor for the lack of ethanol-induced craniofacial malformations at 24 hpf, we repeated starting our exposure paradigm at 24 hpf but increased our ethanol exposure concentration from 1% to 1.3%. We did not see any increase in the penetrance and expressivity of craniofacial malformations compared to the 1% exposure dose (Table S2). Combined, these results suggest that mutation in *bmp2b, bmp4* and *bmpr1bb* sensitizes embryos to ethanol induced facial defects when exposed to ethanol from 10-18 hpf.

### Ethanol alters overall shape of the viscerocranium in Bmp mutants

Micrognathia is a hallmark of ethanol exposure in humans, though recent data has shown greater variation in facial shape in FASD (Blanck-Lubarsch et al., 2020; Suttie et al., 2013). To understand the impact of ethanol on viscerocranial shape and size, we undertook as series of quantitative measures to directly assess facial shape changes in zebrafish. To quantify micrognathia-like reductions in jaw size observed in FASD, we dissected and measured the size of the Meckel’s cartilages from untreated *bmp2b^+/-^* and ethanol-treated wild type and *bmp2b^+/-^* larvae. Ethanol-treated *bmp2b^+/-^* larvae displayed a significant reduction in jaw size compared to untreated *bmp2b^+/-^* and ethanol-treated wild type larvae (Fig. 1I, n=29 larvae, both Meckel’s cartilage elements per group (58 total), one-way ANOVA, F=36.85, p<.0001). Beyond jaw defects, we observed malformations in additional viscerocranial cartilage elements, in particular an increase of the angle between the ceratohyal (Ch) elements (Fig. 1B-D compared to F-H). This suggests that embryonic ethanol exposure is disrupting facial shape.

PAE is known to result in general growth retardation and developmental delay in both humans and animal models (Everson & Eberhart, 2023; Popova et al., 2023). To expand our assessment of Bmp-ethanol interactions on facial shape and control for developmental delays, we performed morphometric analysis on untreated and ethanol-treated wild type and *bmp4^-/-^* larvae. This approach takes into account changes in size when analyzing facial shape by using by Procrustes superimposition which removes variation in size, position and orientation, key to our analyses in overall facial shape (Goodall, 1991; Klingenberg, 2011). We determined facial shape by labeling each joint in the viscerocranium (Fig. 2A, Alcian Blue-stained image). Principle component analysis (PCA) of facial shape revealed that PC1 represented over 50% of facial variation as a shortening and widening of the viscerocranium, as well as a flattening of the angles of several cartilage elements (Fig. 2A, Principal component 1). PC2 represented nearly 20% of the variation in facial shape, affecting width of the midface (Fig. 2A, Principal component 2). Our dataset shows that the greatest amount of variation occurred in ethanol-treated *bmp4^-/-^*larvae, while the smallest variation occurred in wild type, wild type larvae (Fig. 2A, magenta dashed vs black dashed 95% confidence ellipses). Ethanol-treated wild type and untreated *bmp4^-/-^* larvae displayed similar increases in variation compared to untreated wild type larvae, but less than ethanol-treated *bmp4^-/-^*larvae. The mean of each group, displayed as a solid ellipse centered in the 95% confidence ellipse, shows little overlap between the groups (Fig. 2A). Procrustes ANOVA analysis showed that these shape changes were significant (F=10.37, p<.0001). Combined, our morphometric data set shows that ethanol-treated *bmp4^-/-^* larvae displayed significant variation in facial shape; however, either ethanol or *bmp4^-/-^* alone also increased variation in viscerocranial shape compared to untreated wild type larvae. We did not identify this variation in ethanol or *bmp4^-/-^* alone in our initial visual screens. Overall, this data shows that while both ethanol-treated wild type and untreated *bmp4^-/-^* larvae display greater variation of facial shape than untreated wild type larvae, ethanol-treated *bmp4^-/-^* larvae exhibit the greatest variation in facial shape with significant, quantifiable changes in facial size and shape compared to all other groups. This suggests that ethanol- or mutation-induced craniofacial shape changes are not readily identified in visual screens for gross morphology and that Bmp mutation appears to potentiate ethanol-induced facial shape changes.

**Figure 2.**
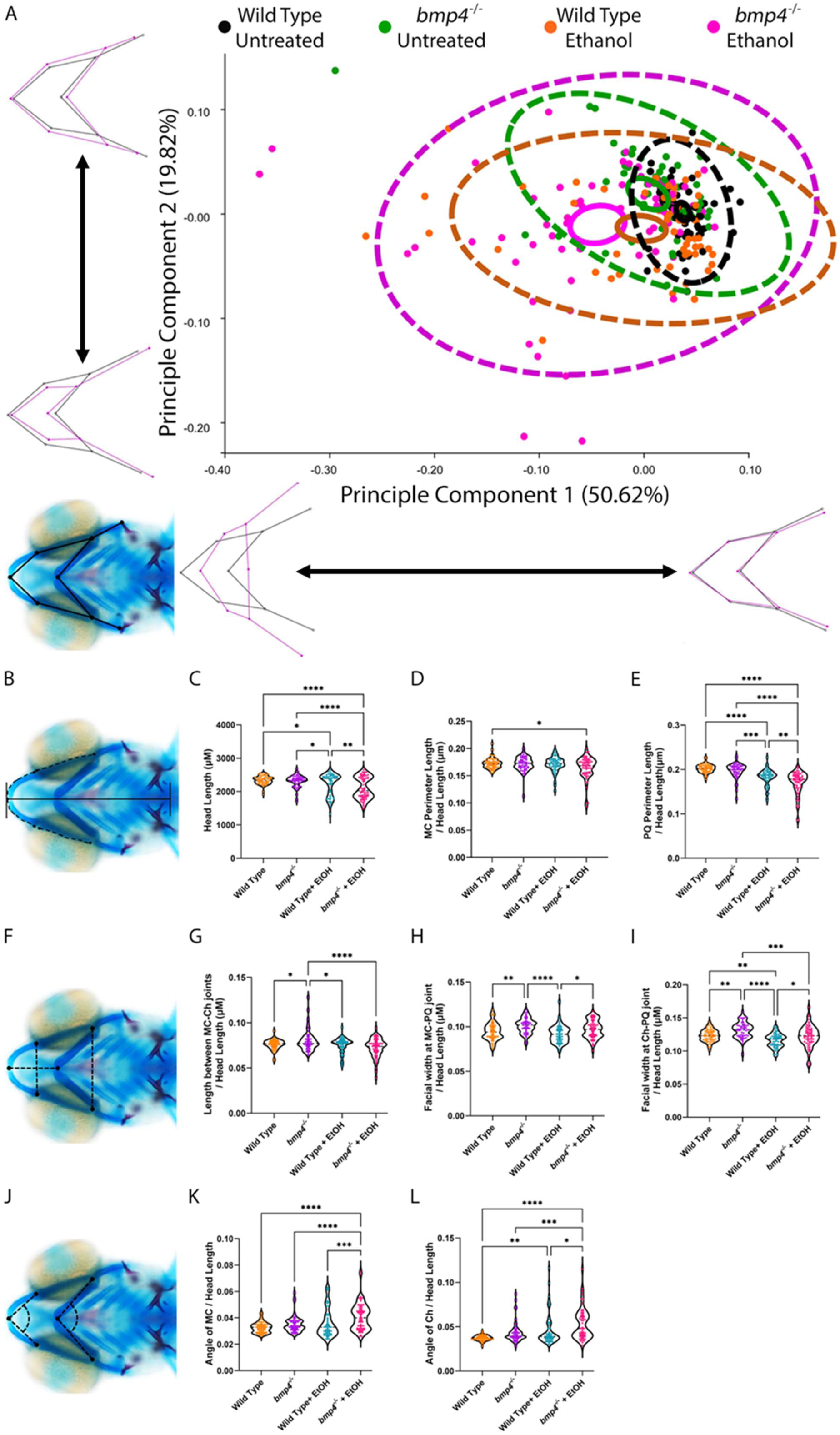
Ethanol exposure alters viscerocranial shape in *bmp4^-/-^* larvae. (**A, B, F**) Whole-mount images of the viscerocranium in 5 dpf larvae showing landmarks, linear measures and cartilage angles (cartilage is blue and bone is red, ventral views, anterior to the left). (**A**) Landmarks were placed on several joints between the cartilage elements of the viscerocranium. Genotypes are color-coded; black = untreated wild type larvae (n = 61), green = untreated *bmp4*^-/-^ larvae (n = 54), orange = ethanol-treated wild type larvae (n = 58), magenta = ethanol-treated *bmp4*^-/-^ larvae (n = 66), dashed circles represent confidence ellipses in which 95% of all individual data points for each group lie, solid circles represent 95% confidence ellipses for means. Wireframe graphs represent variation described by each axis with black representing no variation and magenta representing variation relative to the black wireframe. For example, PC1 captures a shortening and widening in viscerocranial shape, while PC2 represents variation in midfacial width. Procrustes ANOVA showed significant change in the viscerocranial shape (F = 10.37, DF = 36, p = .0001). (**C**) Overall head length. All subsequent measures were analyzed as ratios to overall head length. (**D**) Measures of Meckel’s perimeter length, (**E**) measures of palatoquadrate perimeter length, (**G**) measures between the midline joints of the Meckel’s cartilages and ceratohyal cartilages and width measures between the joints of the (**H**) Meckel’s and palatoquadrate cartilages and (**I)** ceratohyal and palatoquadrate cartilages. Measures of angles joint of the (**K**) Meckel’s and (**L**) ceratohyal cartilages. Linear measures show significant decrease in facial width and length and an increase in angle between cartilage elements, representing flattening of the facial skeleton even when head length is taken into account (Individual graph statistics in Table S3).

To confirm the morphometric data, we performed linear measurements on the viscerocrania of the *bmp4* morphometric dataset. For genotyping we remove the tail of the larvae preventing overall body length measures. However, we measured the overall length of the head as a measure for general developmental delays. We also measured the length between the midline joints of the Meckel’s and ceratohyal cartilages, the width between the joints of the Meckel’s and palatoquadrate cartilages and ceratohyal and palatoquadrate cartilages and the length of the perimeter of the Meckel’s and palatoquadrate cartilages (Fig. 2B & F). We further performed angle measures of the midline joints of the Meckel’s and ceratohyal cartilages to analyze the flattening of the viscerocranium cartilage elements (Fig. 2J). We observed a significant decrease in head length due to ethanol that was further exacerbated by loss of *bmp4* (Fig. 2C, Table S3). Consistent with PC1 (Fig. 2A), we observed a significant decrease in the length between the Meckel’s and ceratohyal cartilages in ethanol-treated *bmp4^-/-^* larvae compared to all other groups (Fig. 2C, Table S3). We observed significant changes in facial width at both the Meckel’s and palatoquadrate cartilages and ceratohyal and palatoquadrate cartilages (Fig. 2D and E, respectively, Table S3). In both measures, ethanol-treated *bmp4^-/-^*larva showed significant decreases in width at both the Meckel’s and palatoquadrate cartilages and ceratohyal and palatoquadrate cartilages compared to untreated wild type larvae and untreated *bmp4^-/-^* larva (Fig. 2D-E, Table S3). In contrast, untreated *bmp4^-/-^* larvae showed an increase in viscerocranial width compared to all other groups; the measurements were significantly increased compared to ethanol-treated wild type and *bmp4^-/-^* larvae (Fig. 2D-E, Table S3) and trending toward significance compared to untreated wild type embryos in Fig. 2D but not in Fig. 2E (Table S3). In angle measures of the Meckel’s and ceratohyal cartilages, ethanol-treated *bmp4^-/-^* larvae showed significant increases in cartilage angles (Fig. 2G-H, Table S3), consistent with the flattening of the viscerocranial cartilages observed initial screens (Fig. 1) and PC1 and PC2 of our morphometric analysis (Fig. 2A). Overall, our morphometric measurements document that the Bmp-ethanol interaction results in a smaller jaw, and shorter and wider viscerocranial shape, consistent with FASD in humans (Bemquerer et al., 2022; Suttie et al., 2013).

### Combinatorial loss of Bmp pathway components exacerbates ethanol-induced viscerocranial malformations

While we consistently observed ethanol-induced viscerocranial shape changes in tested Bmp mutants that differed from wild type controls, both the penetrance and expressivity of these defects were highly variable (Figs. 1-2, Table 1). The Bmp pathway is a complex signaling pathway comprised of multiple components (Little & Mullins, 2009). These components can be partially redundant at different levels of the pathway (Li et al., 2011; Mu et al., 2021; Schille et al., 2016; Wise & Stock, 2010). Given this redundancy, we hypothesized that combinatorial loss of pathway components potentially increases both the penetrance and expressivity of ethanol-induced viscerocranial defects. Using a hypomorphic *smad5* allele that has a highly stereotyped phenotype of cartilage fusions and malformations and have been previously shown to be insensitive to ethanol exposure (McCarthy et al., 2013; Swartz et al., 2011), we generated double-mutant *bmp4^-/-^;smad5^-/-^* larvae (Fig. 3). We found that *bmp4^-/-^;smad5^-/-^* larvae exhibit 100% penetrant, ethanol-malformations that are a) more severe than the stereotypical *smad5* phenotypes in untreated *bmp4^-/-^;smad5^-/-^* induced viscerocranial larvae, b) much more severe than any of the Bmp single mutants (Fig. 3D compared to B; compared to Fig. 1) and c) highly reminiscent of the severe facial phenotypes observed in Dorsomorphin treated embryos (Lovely et al., 2016). We also observed a spectrum of phenotype severity in *bmp4^+/-^;smad5^-/-^* larvae, in which one gene copy of *bmp4* remained wild-type: these phenotypes range from stereotypical *smad5* mutant phenotypes to phenotypes comparable to ethanol-treated double-homozygous *bmp4^-/-^;smad5^-/-^* larvae (Fig. S1). In contrast, we observed no impact of ethanol on double-heterozygous *bmp4^+/-^;smad5^+/-^*larvae (data not shown). These genetic interaction results indicate that the variation in *bmp^+/-^;smad5^-/-^* larvae may be driven either by expression differences in other components of the Bmp pathway or additional yet identified ethanol-sensitive alleles. Ultimately, these data support our previous observation that loss of *bmp4* potentiates ethanol-induced facial shape changes with the most significant impact observed in the combination of mutation and ethanol exposure.

**Figure 3.**
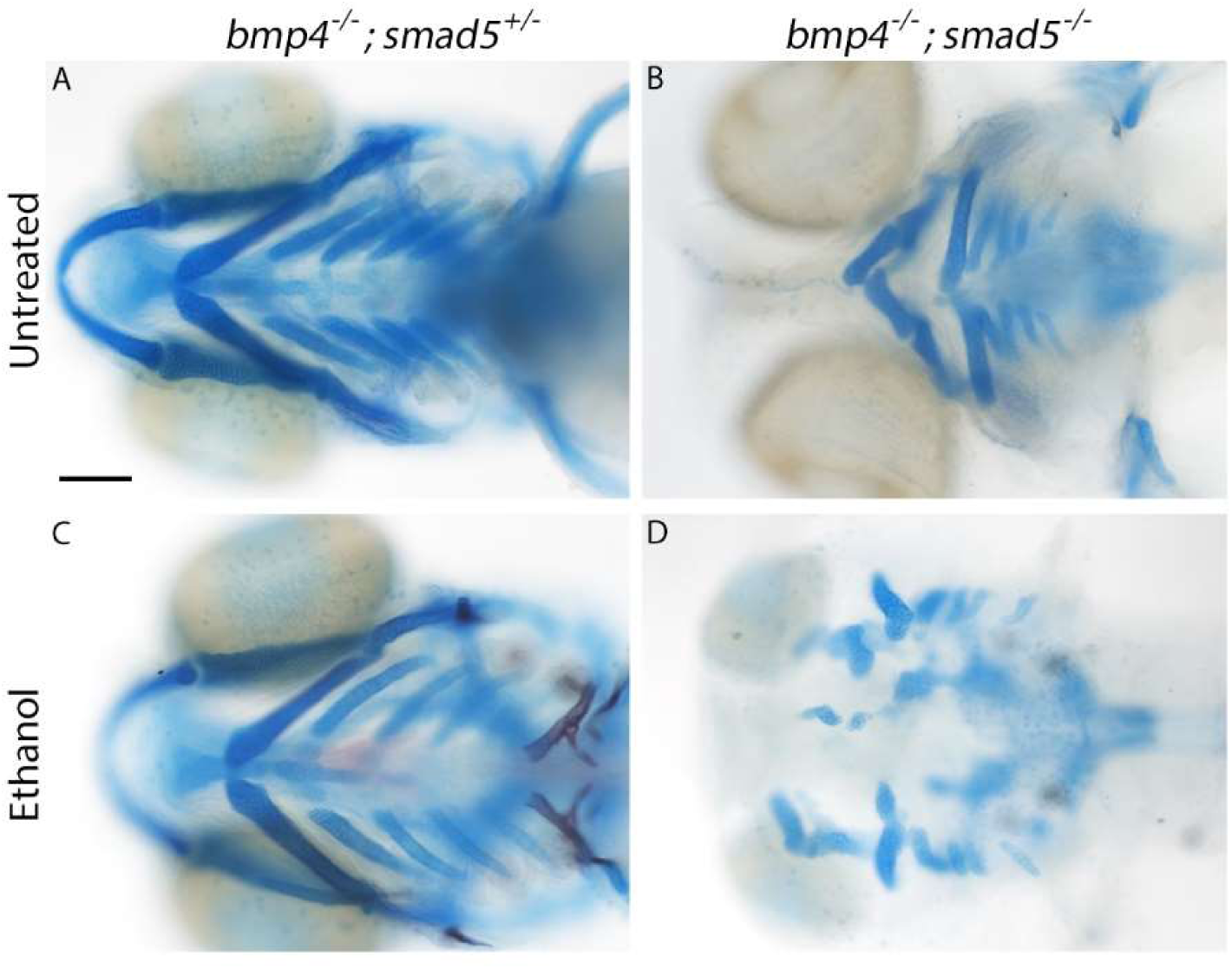
*bmp4^-/-^;smad5^-/-^* larvae exhibit full penetrant, exacerbated ethanol-induced facial malformations. (**A-D**) Whole-mount images of viscerocranium at 5 dpf larvae. Cartilage is blue and bone is red (Ventral views, anterior to the left, scale bar: 100 μm). (**A**) No impact on facial formation was observed in untreated *bmp4^-/-^;smad5^+/-^* larvae. (**B**) Stereotypical *smad5* mutant phenotypes were observed in *bmp4^-/-^;smad5^-/-^* larvae. (**C**) Ethanol does not impact viscerocranial morphology in *bmp4^-/-^;smad5^+/-^* larvae. (**D**) However, *bmp4^-/-^;smad5^-/-^* larvae are fully penetrant, ethanol sensitive, resulting several viscerocranial malformations recapitulating the severe phenotypes in Dorsomorphin-treated embryos (Lovely et al., 2016).

### Ethanol-treated Bmp mutants display malformations to the anterior pharyngeal endoderm and *fgf8a* expression in the oral ectoderm

We have previously shown that zebrafish embryos treated with the Bmp signaling inhibitor compound Dorsomorphin from 10-18 hpf exhibit disrupted morphogenesis of the anterior pharyngeal endoderm (Lovely et al., 2016). Multiple studies have shown that disruptions to the anterior pharyngeal endoderm leads to jaw malformations (Balczerski et al., 2012; Couly et al., 2002; Crump, 2004; Haworth et al., 2004, 2007). To analyze anterior pharyngeal endoderm shape, we labeled the pharyngeal endoderm by crossing the *sox17:EGFP* transgene, which labels all endoderm with EGFP (Chung & Stainier, 2008), into the *bmp4^-/-^;smad5^-/-^* background. Compared to wild type, ethanol-treated *bmp4^-/-^;smad5^-/-^* embryos showed mild changes to overall endoderm shape (Fig. 4B-E). However, our more detailed examination showed that the shape of the anterior pharyngeal endoderm is altered in ethanol-treated *bmp4^-/-^;smad5^-/-^* embryos (Fig. 4B’-E’). To quantify these changes in anterior pharyngeal endoderm shape, we measured width at the first pouch (AE width), midline length (AE length) and area anterior pharyngeal endoderm (AE area) from the first pouch to the anterior most end of the pharyngeal endoderm (Fig. 4F). We normalized for ethanol-induced general growth deficits to anterior pharyngeal endoderm shape, by measuring the length and width of the embryonic head and calculating head area (Fig. 4G). Calculating head area controls for ethanol-induced reductions in eye size and did not show any differences between groups (Fig. 4H). Our analyses showed a significant increase in AE area expressed as a ratio to head area in ethanol-treated double-homozygous *bmp4^-/-^;smad5^-/-^* embryos compared to all other groups (Fig. 4H, Table S4). AE area alone of ethanol-treated *bmp4^-/-^;smad5^-/-^* embryos only exhibited a significant increase compared to ethanol-treated wild type, wild type embryos (Fig. 4I, Table S4). We also observed non-significant increases in AE length and AE width expressed as a ratio to head area in ethanol-treated *bmp4^-/-^;smad5^-/-^* embryos while no differences were observed in head length and width (Fig. S3, Table S5). Collectively, our observations in zebrafish document that changes in anterior pharyngeal endoderm area in ethanol-treated Bmp mutants might underlie viscerocranial malformations.

**Figure 4.**
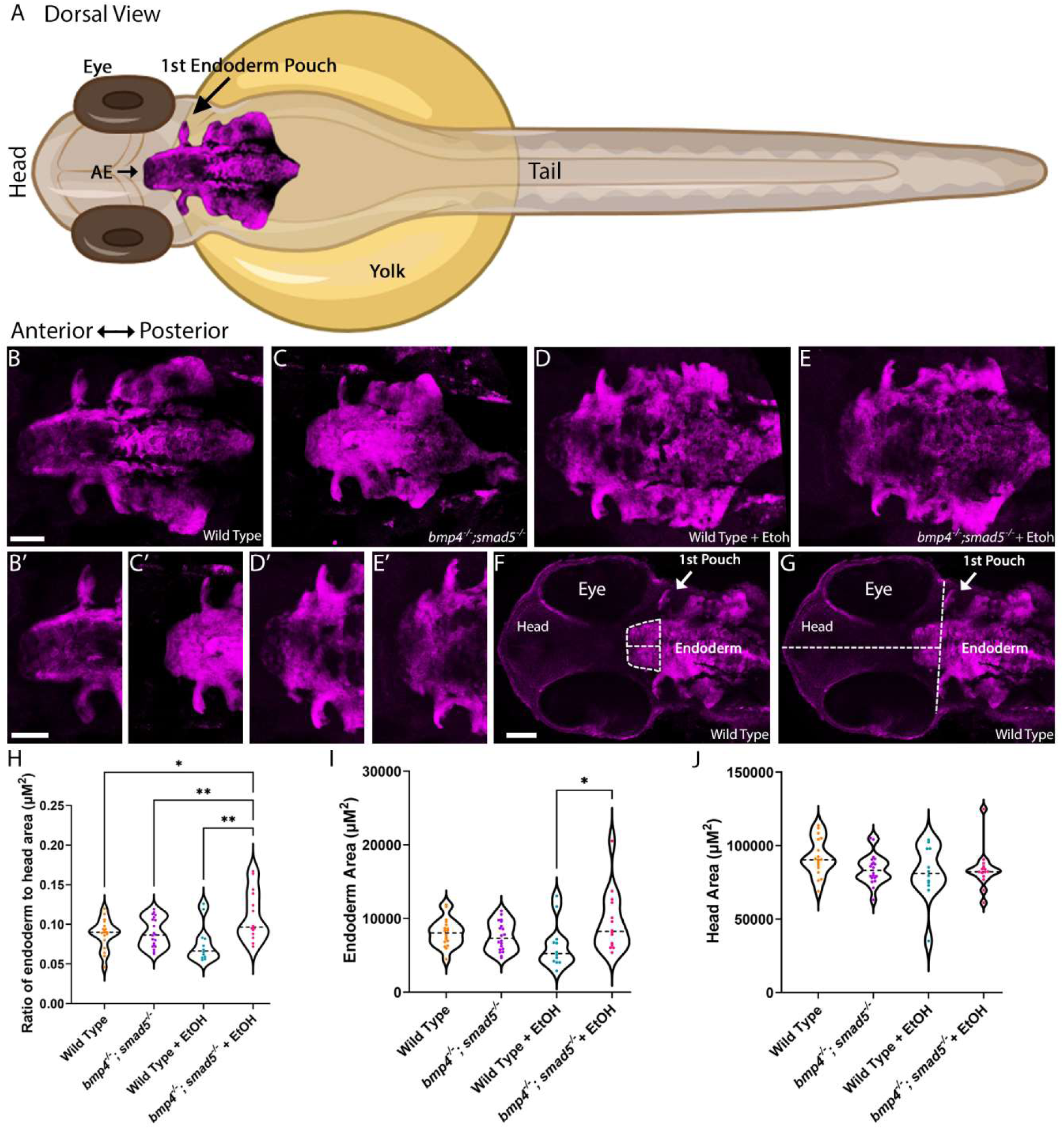
Ethanol exposure alters area of the anterior pharyngeal endoderm in *bmp4^-/-^;smad5^-/-^* embryos. (**A**) Schematic with annotation of the pharyngeal endoderm in a zebrafish embryo (zebrafish image generated from BioRender; AE = Anterior Endoderm). (**B-E**) Whole-mount images of pharyngeal endoderm in 36 hpf embryos; (**B’-E’**) magnified view of the anterior pharyngeal endoderm in (**B, B’)** untreated wild type embryos, n = 18; (**C, C’)** untreated *bmp4^-/-^;smad5^-/-^* embryos, n = 20; (**D, D’)** ethanol-treated wild type embryos, n = 12; (**E,E’)** ethanol-treated *bmp4^-/-^;smad5^-/-^*embryos, n = 14 (dorsal views, anterior to the left; Size bars: 50 μm). (**F**) Area and length of the anterior pharyngeal endoderm was measured from the first pouch to the anterior-most tip while width of the anterior pharyngeal endoderm was measured at the level of the first pouch. (**G**) Area of the head was calculated from length (first pouch to most anterior tip of the head) and width (measured at the level of the first pouch) of the head. (**H**) Ratio of anterior pharyngeal endoderm to head area shows that ethanol-treated *bmp4^-/-^;smad5^-/-^* embryos show increased area of the anterior endoderm compared to all other groups. This increased size is not due to changes in head size (**I**) but due to increased size in the anterior pharyngeal endoderm directly (**J**, individual graph statistics in Table S4).

Previous work has shown that the anterior pharyngeal endoderm is necessary to induce signaling factor expression in the oral ectoderm (Balczerski et al., 2012; Haworth et al., 2004, 2007). This suggests that blocking Bmp signaling with either Dorsomorphin or ethanol will disrupt expression of the critical oral ectoderm markers, such as *fgf8a* and *pdgfaa.* To test if expression of *fgf8a* or *pdgfaa* is altered in Dorsomorphin-treated embryos, we performed HCR-based, fluorescent *in situ* hybridization. In wild type embryos, *fgf8a* is expressed in a small domain in the oral ectoderm, ventro-posterior to the developing eye and retina (Fig. S3A). Wild type embryos treated with Dorsomorphin from 10-18 hpf, the same developmental time window as our ethanol exposure paradigm, have variable defects in *fgf8a* expression from loss of expression to anterior expansion of expression to the level of developing the retina (Fig. S3B & C). This mirrors *fgf8a* expression changes in other anterior pharyngeal endoderm mutants (Balczerski et al., 2012; Haworth et al., 2007). Interestingly, *pdgfaa* which is expressed throughout the oral ectoderm, is unaltered in Dorsomorphin-treated embryos (Fig. S3D & E), demonstrating that the oral ectoderm is not lost when Bmp signaling is attenuated and that the endoderm mediates the expression of some, but not all, oral ectodermal signaling molecules. We then tested if of *fgf8a* is altered in ethanol-treated *bmp4^-/-^* and *bmp4^-/-^;smad5^-/-^* embryos. The expression domain of *fgf8a* was similarly well-detectable in untreated *bmp4^-/-^*, *bmp4^-/-^;smad5^-/-^* and ethanol-treated wild type embryos with embryos from each genotype showing little difference to untreated wild type embryos (Fig. S4A-C; Fig. 5A-C). Expression of *fgf8a* in ethanol-treated *bmp4^-/-^* embryos are subtly expanded anteriorly to the level of developing the retina while *fgf8a* expression *bmp4^-/-^;smad5^-/-^* embryos was markedly expanded anteriorly to the level of developing the retina (Fig. S4D; Fig. 5D). This anterior expansion of the *fgf8a* expression domain is similar to the anterior expansion of *fgf8a* we observed in Dorsomorphin-treated embryos (Fig. S4D; Fig. 5D compared to Fig. S3C). Our data of ethanol-induced changes upon Bmp signaling perturbation to the *fgf8a* expression domain in the oral ectoderm is consistent with previous work showing that malformations of the anterior pharyngeal endoderm disrupting oral ectoderm expression domains and subsequent viscerocranial malformations (Balczerski et al., 2012). While limited to *fgf8* as critical marker, these observations indicate that synergistic Bmp-ethanol perturbations disrupt an anterior pharyngeal endoderm-oral ectoderm-signaling axis.

**Figure 5.**
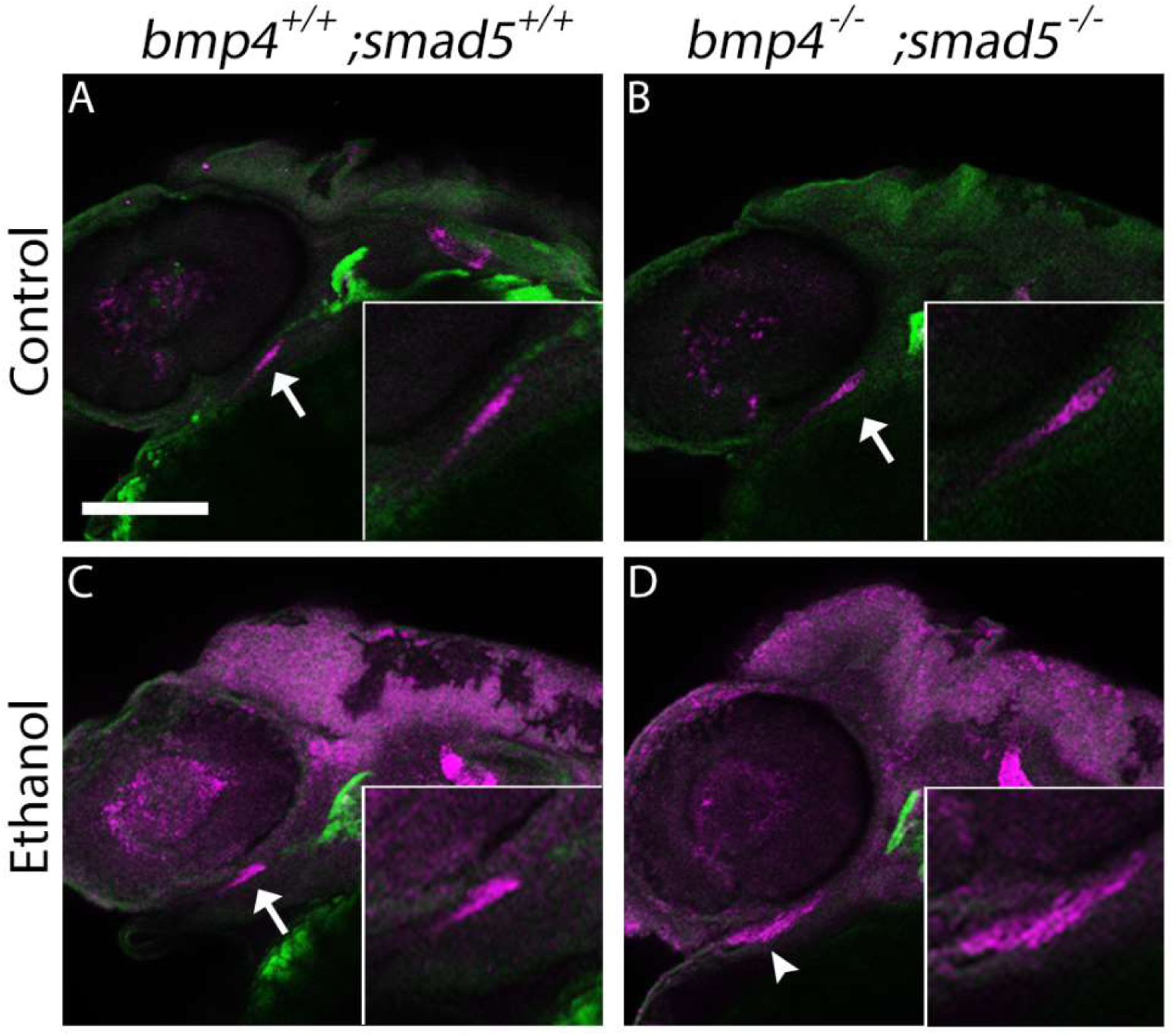
Ethanol exposure changes shape of oral ectoderm expression domain in *bmp4^-/-^;smad5^-/-^* embryos. (**A-D**) Whole-mount, confocal images of *bmp4;smad5;sox17:EGFP* embryos fluorescently labeling *fgf8a* gene expression at 36 hpf (lateral views, anterior to the left, scale bar: 100 μm). (**A-C**) Arrows show normal expression of *fgf8a* in the oral ectoderm of untreated wild type and *bmp4^-/-^;smad5^-/-^* embryos as well as ethanol-treated wild type embryos. (**D**) Arrowhead shows that domain of *fgf8a* expression in ethanol-treated *bmp4^-/-^;smad5^-/-^* embryos is expanded anteriorly (n = 7 embryos per group).

Our data described above suggests that ethanol will reduce Bmp signaling responses in the anterior endoderm. To test this we, generated *bmp4;smad5;sox17GFP;BRE:mKO2,* double mutant, double transgenic line that labels the endoderm in GFP and active Bmp signaling in mKO2 (Collery & Link, 2011). We have previously used this *BRE:mKO2* line to analyze active Bmp response during early pharyngeal arch development (Sheehan-Rooney et al., 2013). Using this line, we show that Bmp signaling is lost in pharyngeal endoderm in *bmp4^-/-^;smad5^-/-^*embryos, independent of ethanol exposure at 18 hpf, the end of Bmp signaling responses in the endoderm (Lovely et al., 2016) (Fig. 6B,B’ & D, D’). However, we did not observe any decrease in the BRE response due to ethanol exposure in other pharyngeal tissues (Fig. 6A-D). We quantified BRE fluorescent levels by measuring the Corrected Total Fluorescence of the BRE response in the pharyngeal arches and observed no significant changes in response to either *bmp4^-/-^;smad5^-/-^* embryos or ethanol (Fig. S5, Table S6). This suggests that ethanol does not impact Bmp signaling but other targets in endoderm morphogenesis.

**Figure 6.**
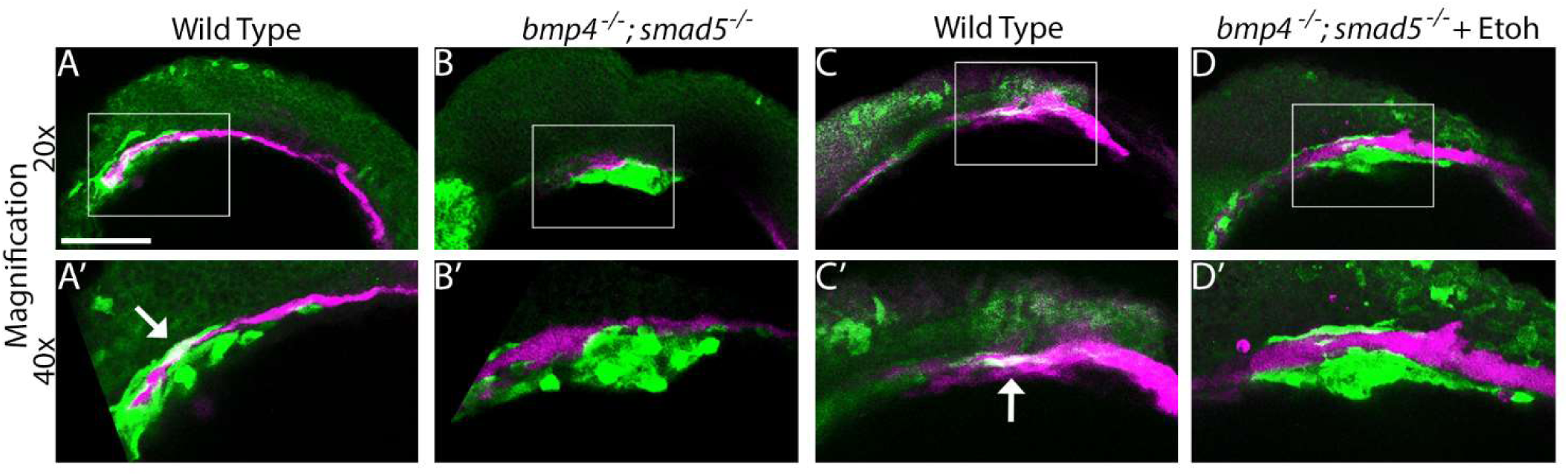
Endoderm-specific Bmp signaling responses are lost in *bmp4^-/-^;smad5^-/-^* embryos but not in ethanol-treated embryos. (**A-D**) Whole-mount, confocal images of *bmp4;smad5;sox17:EGFP;BRE:mKO2* embryos fluorescently labeling the endoderm in GFP (false colored to magenta) and active Bmp signaling with mKO2 (false colored to GFP) at 36 hpf (lateral views, anterior to the left, boxes represent area imaged at 40x, scale bar: 100 μm). (**A,A’ & C,C’**) Arrows show overlap of Bmp response and *sox17* labeled endoderm. (**B,B’ & D,D’**) Endoderm-specific Bmp signaling responses are lost in *bmp4^-/-^;smad5^-/-^* embryos (**B,B’ & D,D’** compared to **A,A’ & C,C’**). Ethanol exposure does not alter Bmp responses (Embryos per group, wild type, n = 5; *bmp4^-/-^;smad5^-/-^*, n = 4; wild type + Etoh, n = 4; *bmp4^-/-^;smad5^-/-^*, n = 8).

### Bmp-ethanol interactions translate to human FASD jaw volume

Zebrafish has been used successfully to model human developmental disorders (Lieschke & Currie, 2007; Santoriello & Zon, 2012) and we have previously shown that our zebrafish screens can predict gene-ethanol interactions in humans (McCarthy et al., 2013). We therefore sought to validate if our zebrafish model of Bmp-ethanol interactions impacting facial shape may be predictive of gene-ethanol associations in humans. From a study of children with and without PAE, we ran a genome-wide association study to explore if genotype was associated with several facial features, including anomalous mandible volume, in the presence or absence of PAE (genotype x PAE interaction). This sample and the genotype x PAE results for various phenotypes are utilized only as a “look-up” resource due to the small sample size (total N=369) and not as a discovery sample (Dou et al., 2018; Hwang et al., 2024; McCarthy et al., 2013) (McCarthy et al,; Dou et al, 2018; Hwang et al, 2024). We examined results for common variants in human *BMP2*, *BMP4*, and *BMPR1B*. Only one SNP in the 5’ region of *BMP2* (*rs235710*) was significant (p=0.029); no other SNPs in or near *BMP2* were correlated or demonstrated association (all p>0.74). Similarly, one SNP in the 5’ region of *BMP4* (*rs72680543*, p=0.043) was significant but was in complete linkage disequilibrium (LD) (D’=1) with two SNPs that were non-significant (*rs72680541*, p=0.19; *rs10873077*, p=0.072). Given the lack of supporting evidence, we did not follow up on the findings in these two genes. The best p-value for SNPs in the BMP receptor gene *BMPR1B* was for *rs34063820* (p=4x10^-4^, Minor Allele Frequency (MAF)=0.12; purple diamond in Fig. 7A).

**Figure 7.**
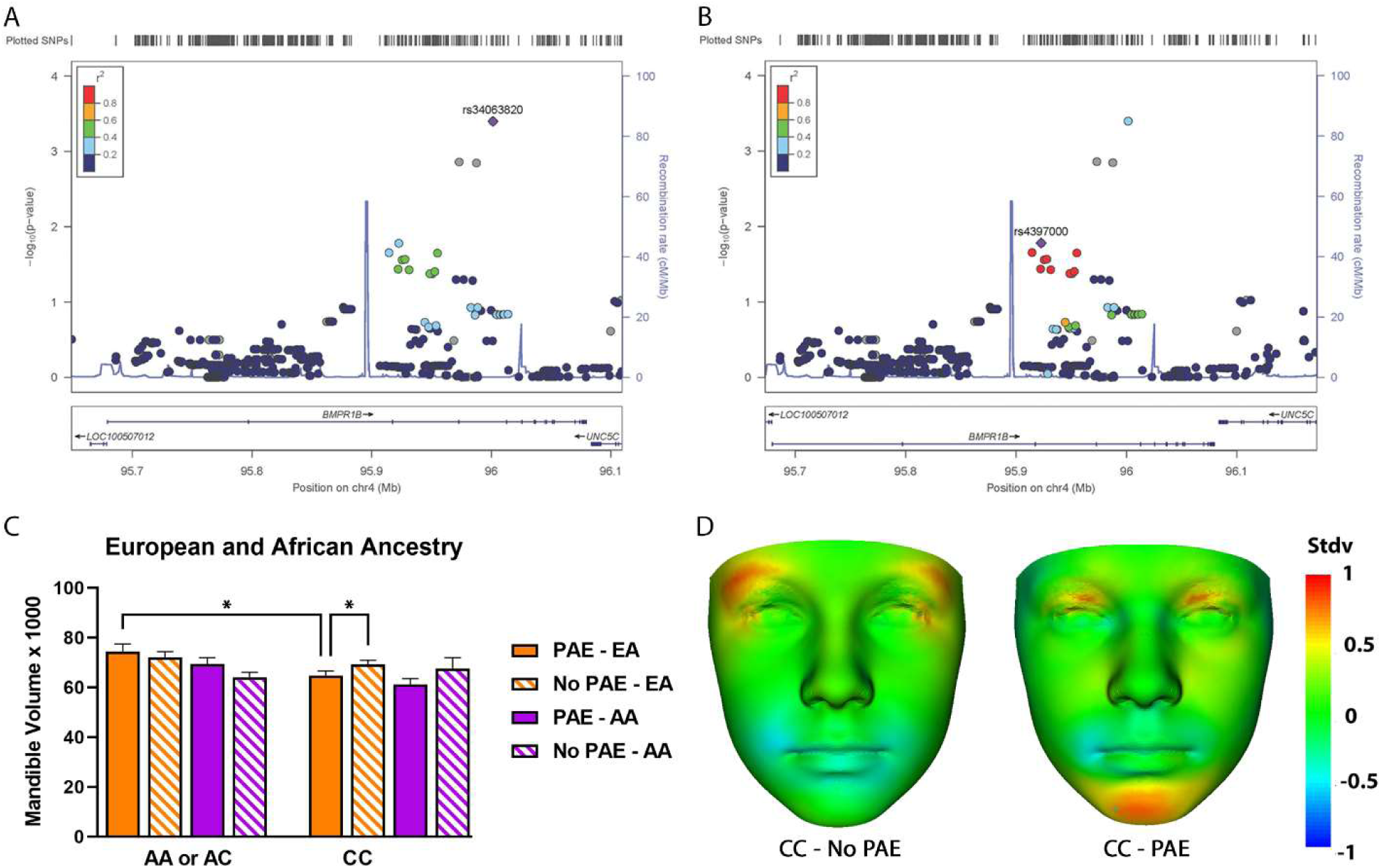
Association of jaw deformations with SNPs located between BMPR1B in a combined European and African American discovery sample. (**A-B**) Y-axis denotes the -log10 (p-value) for the genotype x PAE effect. X-axis is the physical position on the chromosome (Mb). The extent of linkage disequilibrium (as measured by r2) in the 1000 Genomes European reference panel between each SNP (circles) and the purple diamond SNP is indicated by the color scale at the top left. Larger values of r2 indicate greater linkage disequilibrium. (**A**) The SNP with the highest p-value in *BMPR1B*, *rs3406382* (p=4x10^-4^), is labeled with a purple diamond. (**B**) The cluster of nine SNPs in high LD with the purple diamond SNP *rs4397000,* p=0.016, and in lower LD with *rs3406382* (blue circle). SNP = single nucleotide polymorphism, PAE = prenatal alcohol exposure, LD = linkage disequilibrium. (**C**) Mandible volume in children of European and African ancestry (EA and AA, respectively) recruited with and without alcohol exposure according to genotype. In EA individuals with the *CC* genotype, mandible volume was significantly decreased in alcohol exposed vs unexposed individuals. In alcohol exposed EA individual’s mandible volume was decreased in those with the *CC* genotype compared to those with the *AA+AC* genotypes. (**D**) DSM analysis facial signature heatmaps indicating surface to normal displacement at +/-1 s.d. for mean EA individuals. Where red-blue-green coloring indicates a reduction-expansion or agreement with the group being normalized against. (Left) *CC* genotype no PAE (n=52) normalized against *AA+AC* genotype no PAE (n=43). (Right) *CC* genotype with PAE (n=57), normalized against *AA+AC* genotype with PAE (n=28). Red coloring on the mandible’s tip indicates a mandibular retraction or micrognathia. Mandible reduction is not observed without PAE. However, we do observe a red signal on the outer canthi, indicating a reduction in this region in the *CC* genotype group.

There were no other significant SNPs that were highly correlated with *rs34063820* to support this association, as indicated by the lack of red or yellow circles in Figure 7A. As the lack of support could indicate a potential false positive result, we focused on a cluster of nine SNPs, which demonstrated slightly less significant association with mandible volume, and were in modest LD with *rs34063820* (r2<0.60; cluster of blue and green circles in Figure 7A). These nine SNPs were highly correlated with each other, and had similar, significant p-values (the same cluster of SNPs are now red circles in Figure 7B). The strongest genotype by PAE interaction in these nine SNPs was for *rs4397000* (purple diamond in Fig. 10B, p=0.016, all nine SNPs p<0.037). The *rs4397000* SNP had a minor allele (*A>C* allele) frequency of 0.209 in the European Ancestry (EA) sample and 0.496 in the African Ancestry (AA) sample. Examining these SNPs in GTEx revealed that eight of the nine SNPs are eQTLs, expressing *BMPR1B* in several tissues. The minor allele *A* of *rs4397000* is associated with increased *BMPR1B* gene expression in the adrenal gland and decreased expression in the brain, heart, and pancreas. Wilcoxon test revealed that among children with an *AA* or *AC* genotype, there was no difference in mandible volume between the PAE (EA n=29, AA n=38) and no PAE (EA n=41, AA n=62; p=0.33) groups, though mandible volume was trending toward significance in the AA samples (EA p=0.70, AA p=0.052; all comparisons are shown in Fig. 7C). However, in EA individuals with two copies of the *C* allele, those without PAE (EA n=52, AA n=15; Fig. 7C) had larger mandible volume on average than those with PAE (EA n=64, p=0.040; AA n=19, p=0.53). Furthermore, EA individuals with PAE with an *AA* or *AC* genotype had larger mandible volume than those who were *CC* (EA p=0.0058, AA p=0.108; Fig. 7C). A facial signature heatmap on the *CC* group normalized against the *AA* or *AC* group visualizes this volumetric reduction only when PAE is present, which is localized to the most anterior part of the mandible (Fig. 7D). Taken together, this genotype correlation with *BMPR1B* variants suggests that gene-ethanol interactions in the Bmp pathway potentially underlie abnormal jaw size in FASD, underlining the predictive value of our mechanistic studies underlying FASD etiology in zebrafish. However, due to the small sample size, these results could be false positives, and need to be replicated with a larger, independent sample.

## Discussion

Formation of the facial skeleton is driven by a complex, highly coordinated, three-dimensional process involving multiple tissues in the developing head (Knight & Schilling, 2006; Medeiros & Crump, 2012; Murillo-Rincón & Kaucka, 2020). Proper regulation of the cell movements and tissue dynamics within a single tissue and between different tissues are critical for these events (Knight & Schilling, 2006; Medeiros & Crump, 2012; Murillo-Rincón & Kaucka, 2020). These cell behaviors are regulated by numerous signaling pathways, which can be attenuated either genetically and/or environmentally, leading to a cascading effect that disrupts craniofacial development (Knight & Schilling, 2006; Medeiros & Crump, 2012; Murillo-Rincón & Kaucka, 2020). Prenatal ethanol exposure, defined as FASD, results in a highly variable set of phenotypes in the facial skeleton, including the jaw (Blanck-Lubarsch et al., 2020). Genetic risk factors are major drivers of FASD symptomology, providing insight into the cellular and molecular processes potentially disrupted in FASD (Lovely, 2020).

### Bmp-ethanol interactions

Here, we show that mutation in several components of the Bmp pathway sensitize zebrafish embryos to ethanol induced malformations to the viscerocranium. We found that when exposed to ethanol from 10-18 hpf, mutations in *bmp2b*, *bmp4* and *bmpr1bb*, sensitize embryos to a range of ethanol-induced defects to Meckel’s cartilages with both later exposure start points and high doses of ethanol at these later time points resulting in less ethanol-induced facial defects. These defects spanned outright loss of cartilage elements to reductions in cartilage size and were consistent amongst all three Bmp mutant lines. Using a morphometric approach, we observed broad changes to facial shape in ethanol-treated Bmp mutants. Ethanol-treated larva displayed a shorter and wider face relative to untreated controls and a flattening of the angles of several cartilage elements. Much of this variation was not identified in our initial screens, supporting that this approach will identify subtle phenotypes impacting facial shape that would be missed if simply using linear measures of cartilage size (McCarthy et al., 2013; Swartz et al., 2014, 2020). However, the variation of phenotypes observed needs a much greater investigation to address the wide array of facial defects seen. Though not examined, heterozygous *bmp4* and *bmpr1bb* larvae will be critical in shedding light on the complex relationship between ethanol and genotype. This combined with expanded timing and dosage analyses will provide much better insight into the broader phenotypic mechanism occurring in these ethanol-sensitive allele. Equally Important, these broad ethanol-induced facial shape changes in zebrafish are consistent with analyses in human patients where alcohol exposed individuals showed broad and varied facial shaped changes compared to age-matched controls (Bemquerer et al., 2022; Suttie et al., 2013). Overall, our data documents that our morphometric analysis in zebrafish improve the rigor of identifying and quantifying ethanol-induced subtle facial shape changes, modeling the highly variable ethanol-induced facial shape changes in human studies.

Our results propose that ethanol increases craniofacial variation in wild type larvae and loss of *bmp4* potentiates this interaction. Exacerbation of facial phenotypes in *bmp4^-/-^;smad5^-/-^*larvae demonstrates that dose-dependent reduction in Bmp signaling drives ethanol-induced phenotypes. Surprisingly, ethanol did not impact Bmp signaling responses, albeit we observed subtle non-significant increases in Bmp signaling responses at 18 hpf. This result was unexpected based on our facial and endoderm analyses as we expected Bmp signaling to be reduced due to ethanol exposure. It is possible that we need to examine earlier time windows as 18 hpf may be to late to observed ethanol-induced changes to Bmp signaling. It is also possible that the loss of Bmp signaling responses the endoderm mask any impact of ethanol on Bmp signaling. A counter hypothesis is that ethanol does not impact Bmp signaling in any meaningful way during pharyngeal development and that ethanol acts on addition targets that regulate some aspect of endoderm morphogenesis and/or craniofacial development. In this case, ethanol would act either downstream of Bmp signaling impinging on target gene expression/function or on a parallel pathway, independent of, and concomitant with Bmp signaling.

Ethanol can impact a number of epigenetic mechanisms, including chromatin modifications (Zakhari, 2013). Recent work has shown that the protein arginine methyltransferase PRMT1 can mediate Bmp-dependent Smad phosphorylation and DNA methylation regulating bone formation and suture closure in mice and ethanol has been shown to regulate both PRMT expression and function (Hashimoto et al., 2017; Ye et al., 2023; Zhao et al., 2019). This suggests that these epigenetic changes can disrupt the expression of Bmp target genes directly, modulating pathway function and target gene expression, or indirectly through methylation patterns that target expression of Bmp components and/or downstream targets. Additionally, these gene expression changes could extend beyond the Bmp pathway and its downstream targets, affecting parallel pathways necessary for craniofacial development. Ultimately, a comprehensive approach combining transcriptomic, proteomic and metabolomic methodologies investigating downstream or parallel pathway targets will be needed to further elucidate the observed variation in the Bmp-ethanol interactions during jaw formation.

### Impact of ethanol on pharyngeal endoderm morphogenesis

Work in multiple model systems has shown that disruption to pharyngeal endoderm morphogenesis leads to jaw defects (Balczerski et al., 2012; Couly et al., 2002; Crump, 2004; Haworth et al., 2004, 2007). This disruption alters the endodermal signaling centers that directly pattern the jaw leading to jaw hypoplasia (Couly et al., 2002; Suzuki et al., 1999; Vieux-Rochas et al., 2010). These signaling centers are critical for neural crest cell survival with disruptions to endoderm morphogenesis leading to neural crest cell death (Edlund et al., 2014; Johnson et al., 2011; Kopinke et al., 2006). We have previously shown that Bmp signaling is required for pharyngeal endoderm morphogenesis from 10-18 hpf, the same as our ethanol-sensitive time window, and blocking Bmp signaling with Dorsomorphin during this time window results in a wide range of defects to the viscerocranium (Lovely et al., 2016). In addition, we observed an increase in cell death in tissues adjacent to the pharyngeal endoderm where neural crest populate suggesting that the Bmp-induced defects in endoderm morphogenesis are increasing cell death in the neural crest (Lovely et al., 2016). Here, we show that our Bmp-ethanol interactions alter anterior pharyngeal endoderm shape, increasing the area of the anterior pharyngeal endoderm relative to head size. Strikingly, this increase in anterior pharyngeal endoderm size stands at odds with the decreases we observed in jaw shape and size. This has been previously observed in morpholino knockdown of *vgll2a*, which results in shorter and wider endodermal pouches but viscerocranial hypoplasia as a result of increased neural crest cell death (Johnson et al., 2011). It is possible that ethanol-induced increases in anterior endoderm size disrupt endodermal signaling centers leading to neural crest cell death. This results in decreased neural crest cell contribution to the forming jaw decreasing its size. However, future work examining endoderm-induced neural crest cell death will need to be properly controlled as ethanol exposure is known to induce neural crest cell death directly (Dunty et al., 2001; McCarthy et al., 2013).

While the anterior pharyngeal endoderm signals directly to the cranial neural crest for cell survival, the signaling can also be indirect, by inducing oral ectoderm signaling factor expression which is critical for jaw formation (Balczerski et al., 2012; Haworth et al., 2004, 2007). These tripartite tissue interactions can be highly variable with endoderm mutants showing both reductions and expansion in oral ectoderm signaling centers and resultant hyper- and hypoplasia of the viscerocranium (Balczerski et al., 2012). Here, we show that, in addition to endodermal defects, blocking Bmp signaling with Dorsomorphin leads to variable disruption of *fgf8a* expression in the oral ectoderm. Strikingly, expression of *fgf8a* in the oral ectoderm was expanded in ethanol-treated *bmp4^-/-^;smad5^-/-^*embryos. Though consistent with the increase in anterior pharyngeal endoderm size, this stands at odds with disruption of Bmp signaling. This suggests that the Bmp-ethanol interaction disrupts jaw formation through an endoderm-oral ectoderm-neural crest signaling axis. However, how this increase in size in both anterior pharyngeal endoderm and oral ectoderm signaling centers differs from Dorsomorphin blockade of Bmp signaling and still results in a smaller jaw in ethanol-treated Bmp mutants remains unclear. Both, the oral ectoderm and surrounding neural crest express a number of signaling factors that drive jaw and palate formation (Swartz et al., 2011). Changes in the expression domain of any number of these could disrupt local interaction points between the neural crest and oral ectoderm resulting in jaw malformations (Balczerski et al., 2012). Of great interest would be the effect of these early developmental disruptions on subsequent cartilage outgrowth, raising several questions cellular events of jaw outgrowth downstream of these tripartite interactions. While our data supports the hypothesis that Bmp-ethanol interactions disrupt an endoderm-oral ectoderm-neural crest-signaling axis, more work needs to be done to analyze the impact of ethanol on the cell behaviors that establish these tripartite interactions and how these interactions drive the cellular mechanisms in the oral ectoderm and neural crest during subsequent jaw formation.

### Zebrafish model gene-ethanol interactions in human FASD

We have previously shown that our zebrafish screens can predict gene-ethanol interactions in humans suggesting that our Bmp-ethanol interactions impacting facial shape may also be predictive of gene-ethanol associations in humans (McCarthy et al., 2013). Consistent with this, disruption of *Bmp2* (*BMP2*) in both mice and humans, *in the absence of ethanol*, result in a hypoplastic jaw (Chen et al., 2019; Sahoo et al., 2011). From a study of children recruited with and without PAE, we show here that several SNPs in *BMPR1B* are significantly associated with ethanol-associated jaw malformations. However, this could be a false positive result, given the small sample size, and needs to be replicated with a larger independent sample. Jaw hypoplasia is commonly observed in FASD (Basart et al., 2018; Blanck-Lubarsch et al., 2020; Suttie et al., 2013) but studying FASD in humans is incredibly challenging due to the complex interplay between genetic background and ethanol timing and dosage. Our results show that zebrafish analyses can model gene-ethanol associations in humans, strongly phenocopying both the malformation and the variation inherent in human data (McCarthy et al., 2013; Suttie et al., 2013, 2017). However, due to the paucity of human genetic studies of FASD, sample sizes with sufficient power to detect association due to the effect of multiple genes in one epistatic model is currently not possible. Thus, the zebrafish model remains a powerful, efficient method to simultaneously examine the effect of multiple genes on facial measures and to generate a deeper mechanistic understanding of these gene-ethanol interactions on craniofacial development. While future functional human and zebrafish analyses will need to test for the causal relationship and mechanistic underpinnings of ethanol-induced jaw deformations, this work strongly suggests that jaw malformations in FASD may in part be due to disruptions to epithelial dynamics and signaling events.

Combined, our results show that zebrafish can predict Bmp-ethanol associations in human and provide a valuable model system for determining the ethanol-sensitive tissue events that contribute to facial defects in FASD. However, despite our increased understanding of the clinical impact of prenatal ethanol exposure, much remains to be learned of the cellular mechanisms underlying FASD (Lovely, 2020). Our work here provides some of the first evidence of gene-ethanol interactions altering epithelial dynamics in the complex, endoderm-oral ectoderm-neural crest-signaling axis leading to facial malformations. This expands our current understanding of ethanol-sensitive tissue dynamics in FASD and provides a conceptual framework for future FASD studies. Ultimately, this work generates a mechanistic paradigm of ethanol-induced structural birth defects and connects ethanol exposure with concrete cellular events that could be sensitive beyond the jaw.

## Materials and Methods

### Zebrafish (*Danio rerio*) care and use

All zebrafish were raised and cared for using established IACUC protocols approved by the University of Louisville (Westerfield, 2007). Adult fish were maintained at 28.5°C with a 14 / 10-hour light / dark cycle. The *bmp2b^tc300a^* (Mullins et al., 1996), *bmp4^st72^*(Stickney et al., 2007), *bmpr1bb^sw40^* (Neumann et al., 2011) and *smad5^b1100^* (Swartz et al., 2011), *sox17:EGFP^s870^* (Chung & Stainier, 2008) and *BRE:mKO2^mw40^* (Collery & Link, 2011) zebrafish lines were previously described. Sex as a biological variable is not applicable at our studied development stages as sex is first detectable in zebrafish around 20-25 days post-fertilization (Aharon & Marlow, 2022), after all of our analyses.

### Zebrafish staging and ethanol treatment

Eggs from random heterozygous crosses were collected and embryos were morphologically staged (Westerfield, 2007), sorted into sample groups of 100 and reared at 28.5°C to desired developmental time points. All groups were incubated in embryo media (EM). At 10 hpf, EM was changed to either fresh EM or EM containing 1% ethanol (v/v). At 18 hpf, EM containing ethanol was washed out with three fresh changes of EM.

### Hybridization chain reaction (HCR), immunofluorescence and in situ hybridization

Embryos were collected at 36 hpf, dechorionated and fixed in 4% paraformaldehyde/PBS at 4°C. HCR protocol was previously described (Ibarra-García-Padilla et al., 2021). HCR amplifiers and buffers were acquired from Molecular Instruments (Molecular Instruments). HCR probes against *fgf8a* and *pdgfaa* were designed as previously described (Kuehn et al., 2022). Immunofluorescence was performed as previously described (Lovely et al., 2016). Primary antibodies were anti-GFP (1:200, sc-9996, Santa Cruz) (Lovely et al., 2016) and anti-mKO2 (1:200, PM051M, MBL) (Gillotay et al., 2020), secondary antibodies were Alexa Fluor 488 and Alexa Fluor 568 (1:500, A10042 and A21124, Invitrogen) (Lovely et al., 2016).

### Imaging & analysis of anterior pharyngeal endoderm shape and Bmp signaling responses

Confocal images were taken using an Olympus FV1000 microscope and measured in FIJI (Schindelin et al., 2012). The anterior pharyngeal endoderm was defined as the medial endoderm anterior to the first pouch. General head area was defined as the product of the width and length of the embryo anterior to the first pouch. *BRE:mKO2* fluorescent intensity was measured in image J by calculating the integrated density of the Bmp response in the pharyngeal arches as well as an average of three measures of mean background fluorescence (the fluorescence of the black background of the image. From these measures, the Corrected Total Fluorescence (CTF) was calculated as Integrated Density – (Area x Mean background fluorescence).

### Cartilage and Bone staining

Zebrafish larva fixed at 5 dpf were stained with alcian blue for cartilage and alizarin red for bone (Walker & Kimmel, 2007). Whole mount, ventral view, brightfield images of the viscerocranium were taken on an Olympus BX53 compound microscope.

### Morphometric analyses

Morphometric analysis of alcian/alizarin-stained larva was performed in TpsDig2 (https://sbmorphomectrics.org) and MorphoJ (Klingenberg, 2011). Landmarks were placed on the following joints, Meckel’s cartilage midline joint, the joints between Meckel’s’ and the palatoquadrate, the palatoquadrate and ceratohyal and at the end of the hyomandibular cartilages. Linear measures were analyzed using TpsDig2. Principle component analysis (PCA), Procrustes ANOVA and wireframe graphs of facial variation were generated using MorphoJ.

### Statistical analyses

Meckel’s cartilage area was analyzed with a one-way ANOVA with a Tukey’s Multiple Comparisons Test. Area measures of the anterior endoderm/head and linear measures/angles of both the anterior endoderm and Alcian/alizarin-stained viscerocranium were analyzed with a two-way ANOVA (type III) with a Tukey’s Multiple Comparisons Test in Graphpad Prism 9.5.1 (Graphpad Software Inc., La Jolla, CA).

### Human Studies

Human participants were recruited as part of the Collaborative Initiative on FASD (CIFASD) from (2007-2017) from sites in Atlanta, GA, Los Angeles, CA, Minneapolis, MN, and San Diego, CA (Mattson, Foroud, et al., 2010; Mattson, Roesch, et al., 2010). Institutional Review Board approval was obtained at each site. All participants and/or their parents/legal guardians provided written informed consent and Institutional Review Board approval was obtained at each recruiting site. Children (aged 5-18 years) who experienced heavy (>4 drinks/occasion at least weekly or >13 drinks/week) prenatal alcohol exposure (PAE) with or without a diagnosis of FAS were classified as PAE, and those who were exposed to minimal (<1 drink/week or <2 drinks/occasion) or no PAE were classified as no PAE (Mattson, Foroud, et al., 2010; Mattson, Roesch, et al., 2010). 3D images were obtained on most participants at the time of the dysmorphology exam using static stereophotogrammetric camera systems capable of capturing a 180-degree image of facial surface geometry. Images were annotated with a sparse set of 20 reliable anthropometric landmarks (Fig. S6A). We performed a Procrustes alignment on the landmarks of each face, aligning them to a template face using a similarity transform, which was applied to each 3D surface to normalize size to a uniform scale. Dense surface models (DSMs) have been used for the assessment of subtle facial dysmorphia across the FASD spectrum (Suttie et al., 2013, 2017), and provide a method to assess surface-based differences of 3D structures, and compare groups of individuals to assess mean differences. A DSM containing 303 individuals was constructed from the uniformly scaled images to produce shape only morphometric models. DSMs allow us to compute facial signatures, which represent normalized differences between groups, and visualize using a heatmap representation group or individual differences (Hammond et al., 2012). To evaluate micrognathia, we computed mandible volume, which was outlined and validated against CT images in (Basart et al., 2018). This technique estimated the volume of a trapezoid formed by four specific points on the size-normalized face: the left and right lower otobasion inferius (lowest points of the ears), gnathion (tip of the mandible), and lower lip vermillion center (Fig. S6B).

Genome-wide association study (GWAS) data were genotyped on the OmniExpress genome array (Illumina, San Diego CA) and on the Multi-Ethnic Genotyping Array at the Johns Hopkins Genetic Resources Core Facility (Baltimore, MD; Dou et al., 2018). Following a previously published GWAS cleaning pipeline (Schwantes-An et al., 2021), the two datasets were cleaned for sample and variant call rates, Hardy-Weinberg Equilibrium (HWE), sample identity using genetic (calculated from single nucleotide polymorphisms [SNPs] on X and Y chromosomes) and self-reported sex, sample relatedness, and genetic ancestry. The two cleaned datasets were imputed separately using Michigan Imputation Server (Das et al., 2016) to 1000-Genomes Phase 3, b37 reference panel (Fairley et al., 2020) then combined. The final dataset consisted of 4,000,362 genotyped and imputed SNPs with minor allele frequency (MAF) ≥ 0.01, genotype rate ≥0.99, and Hardy-Weinberg equilibrium p≥0.000001. Principal components analysis (PCA) was performed using SNPRelate (Zheng et al., 2012) with genotype data from autosomes. Individuals in the 1000-Genomes database (Fairley et al., 2020) were included as reference samples for clustering individuals based on genetic ancestry similarities. Individuals were grouped with European Ancestry (EA, n=222), African Ancestry (AA, n=103) or other (n=44) ancestry samples from 1000 Genomes Project using the first three principal components. Only EA and AA individuals were included in our analyses.

Analyses were performed separately in the EA and AA groups using R (version 4.2.0; R Foundation for Statistical Computing; http://cran.r-project.org/) and PLINK v2.00a3 64-bit (8 Jun 2021). Association of SNP genotype with mandible volume was assessed using an additive generalized linear model with sex, age at time of 3D image, the first 10 genetic principal components, genotype, PAE and genotype by PAE interaction. As the effect of interest was the association of genotype with mandible volume with and without the presence of PAE, the p-value for the genotype by PAE interaction is reported. Variants within 25 kilobase pairs of the three genes of interest (*BMP2, BMP4, BMPR1B*) were evaluated in the EA group to identify significant SNPs (p<0.05). To further explore the genotype by PAE interaction, we employed a non-parametric Wilcoxon test (SAS v 9.4) to compare mandible volume in individuals with PAE to those without PAE stratified by genotype, and to compare mandible volume by genotype groups stratified by PAE. All analyses were performed in the EA and AA samples separately.

## Acknowledgements

The authors like to thank Kevin Kump for zebrafish animal care and husbandry. We thank Dr. Jim Amatruda for the providing the *bmpr1bb* zebrafish line. We also thank Dr. Duygu Özpolat and Dr. Ryan Hull for providing script for HCR probe design. We thank Dr. Christian Mosimann for external manuscript editing. Data used in preparation of this article were obtained in conjunction with the Collaborative Initiative on Fetal Alcohol Spectrum Disorders (CIFASD; https://cifasd.org/data-sharing/) supported by the National Institute of Alcohol Abuse and Alcoholism.

## Data and materials availability

The human genetic data utilized in the CIFASD cohort are available at https://cifasd.org/data-sharing/.

## Author Contributions

CBL conceived the project. JRK and CBL designed all zebrafish studies. JRK, RG, HV, GC and CBL conducted all zebrafish experiments. JRK and CBL analyzed zebrafish data and generated all zebrafish figures. LW, TS, MA and MS designed all human studies and analyzed human data. LW and MS generated human figures. JLK and CBL wrote the manuscript. LW, MS, TF, JLK and CBL reviewed and edited the manuscript. All authors have read and approved the final manuscript.

---

**Supplemental Table 1.** Timing of *bmp4*-ethanol penetrance. Percent of jaw malformations and jaw loss per exposure start window in ethanol-treated *bmp4* mutant embryos generated from random heterozygous crosses.

| Experiments | Untreated Embryos | EtOH-treated Embryos | Jaw Malformations | Percent | Jaw Loss | Percent |
| --- | --- | --- | --- | --- | --- | --- |
| Control | 97 |  | 0 | 0.0% | 0 | 0.0% |
| E10 |  | 174 | 42 | 24.1% | 6 | 3.4% |
| E14 |  | 193 | 23 | 11.9% | 7 | 3.6% |
| E18 |  | 192 | 18 | 9.4% | 1 | 0.5% |

**Supplemental Table 2.** Dose response of *bmp4*-ethanol penetrance. Percent of jaw malformations and jaw loss per dose at starting at 24 hpf in ethanol-treated *bmp4* mutant embryos generated from random heterozygous crosses.

| Experiments | Untreated Embryos | EtOH-treated Embryos | Jaw Malformations | Percent | Jaw Loss | Percent |
| --- | --- | --- | --- | --- | --- | --- |
| Control | 138 |  | 0 | 0.0% | 0 | 0.0% |
| E 1% |  | 165 | 2 | 1.2% | 0 | 0.0% |
| E 1.1% |  | 159 | 2 | 1.3% | 0 | 0.0% |
| E 1.2% |  | 165 | 2 | 1.2% | 0 | 0.0% |
| E 1.3% |  | 159 | 2 | 1.3% | 0 | 0.0% |

**Supplemental Table 3.** Statistical analyses of linear facial measures in Figure 2. Alcian/alizarin-stained viscerocraniums were analyzed with a two-way ANOVA (type III). F-statistic and P-value for each analysis are shown. A Tukey’s Multiple Comparisons Test for each comparison are shown.

| Length at MC-Ch joints / Head Length |  |  | Width at MC/PQ joint / Head Length |  |  | Width at Ch-PQ joint / Head Length |  |  |
| --- | --- | --- | --- | --- | --- | --- | --- | --- |
| ANOVA | F = 7.689 | P < 0.0001 | ANOVA | F = 9.747 | P < 0.0001 | ANOVA | F = 14.96 | P < 0.0001 |
| Tukey's Comparisons |  |  | Tukey's Comparisons |  |  | Tukey's Comparisons |  |  |
| Wild type vs *bmp4*<sup>-/-</sup> | p = 0.05 |  | Wild type vs *bmp4*<sup>-/-</sup> | p = 0.002 |  | Wild type vs *bmp4*<sup>-/-</sup> | p = 0.0033 |  |
| Wild type vs wild type + EtOH | p = 0.9966 |  | Wild type vs wild type + EtOH | p = 0.3758 |  | Wild type vs wild type + EtOH | p = 0.0073 |  |
| Wild type vs *bmp4*<sup>-/-</sup> + EtOH | p = .1067 |  | Wild type vs *bmp4*<sup>-/-</sup> + EtOH | p = 0.4716 |  | Wild type vs *bmp4*<sup>-/-</sup> + EtOH | p = 0.957 |  |
| *bmp4*<sup>-/-</sup> vs wild type + EtOH | p = 0.0292 |  | *bmp4*<sup>-/-</sup> vs wild type + EtOH | p < 0.0001 |  | *bmp4*<sup>-/-</sup> vs wild type + EtOH | p < 0.0001 |  |
| *bmp4*<sup>-/-</sup> vs *bmp4*<sup>-/-</sup> + EtOH | p < 0.0001 |  | *bmp4*<sup>-/-</sup> vs *bmp4*<sup>-/-</sup> + EtOH | p = .1074 |  | *bmp4*<sup>-/-</sup> vs *bmp4*<sup>-/-</sup> + EtOH | p = 0.0005 |  |
| Wild type + EtOH vs *bmp4*<sup>-/-</sup> + EtOH | p = 0.1838 |  | Wild type + EtOH vs *bmp4*<sup>-/-</sup> + EtOH | p = 0.0145 |  | Wild type + EtOH vs *bmp4*<sup>-/-</sup> + EtOH | p = 0.0305 |  |
| Angle at MC / Head Length |  |  | Angle at Ch / Head Length |  |  | Head Length |  |  |
| ANOVA | F = 15.02 | P < 0.0001 | ANOVA | F = 14.89 | P < 0.0001 | ANOVA | F = 18.46 | P < 0.0001 |
| Tukey's Comparisons |  |  | Tukey's Comparisons |  |  | Tukey's Comparisons |  |  |
| Wild type vs *bmp4*<sup>-/-</sup> | p = 0.3508 |  | Wild type vs *bmp4*<sup>-/-</sup> | p = 0.0674 |  | Wild type vs *bmp4*<sup>-/-</sup> | p = 0.9725 |  |
| Wild type vs wild type + EtOH | p = 0.1032 |  | Wild type vs wild type + EtOH | p = 0.0048 |  | Wild type vs wild type + EtOH | p = 0.011 |  |
| Wild type vs *bmp4*<sup>-/-</sup> + EtOH | p < 0.0001 |  | Wild type vs *bmp4*<sup>-/-</sup> + EtOH | p < 0.0001 |  | Wild type vs *bmp4*<sup>-/-</sup> + EtOH | p < 0.0001 |  |
| *bmp4*<sup>-/-</sup> vs wild type + EtOH | p = 0.9371 |  | *bmp4*<sup>-/-</sup> vs wild type + EtOH | p = 0.8394 |  | *bmp4*<sup>-/-</sup> vs wild type + EtOH | p = 0.0477 |  |
| *bmp4*<sup>-/-</sup> vs *bmp4*<sup>-/-</sup> + EtOH | p < 0.0001 |  | *bmp4*<sup>-/-</sup> vs *bmp4*<sup>-/-</sup> + EtOH | p = 0.001 |  | *bmp4*<sup>-/-</sup> vs *bmp4*<sup>-/-</sup> + EtOH | p < 0.0001 |  |
| Wild type + EtOH vs *bmp4*<sup>-/-</sup> + EtOH | p = 0.0005 |  | Wild type + EtOH vs *bmp4*<sup>-/-</sup> + EtOH | p = 0.0128 |  | Wild type + EtOH vs *bmp4*<sup>-/-</sup> + EtOH | p = 0.0047 |  |
| Perimeter Length of MC |  |  | Perimeter Length of PQ |  |  |  |  |  |
| ANOVA | F = 3.525 | P = 0.0163 | ANOVA | F = 36.27 | P < 0.0001 |  |  |  |
| Tukey's Comparisons |  |  | Tukey's Comparisons |  |  |  |  |  |
| Wild type vs *bmp4*<sup>-/-</sup> | p = 0.9917 |  | Wild type vs *bmp4*<sup>-/-</sup> | p = 0.4074 |  |  |  |  |
| Wild type vs wild type + EtOH | p = 0.9748 |  | Wild type vs wild type + EtOH | p < 0.0001 |  |  |  |  |
| Wild type vs *bmp4*<sup>-/-</sup> + EtOH | p = 0.0223 |  | Wild type vs *bmp4*<sup>-/-</sup> + EtOH | p < 0.0001 |  |  |  |  |
| *bmp4*<sup>-/-</sup> vs wild type + EtOH | p = 0.9993 |  | *bmp4*<sup>-/-</sup> vs wild type + EtOH | p = 0.0002 |  |  |  |  |
| *bmp4*<sup>-/-</sup> vs *bmp4*<sup>-/-</sup> + EtOH | p = 0.067 |  | *bmp4*<sup>-/-</sup> vs *bmp4*<sup>-/-</sup> + EtOH | p < 0.0001 |  |  |  |  |
| Wild type + EtOH vs *bmp4*<sup>-/-</sup> + EtOH | p = 0.0786 |  | Wild type + EtOH vs *bmp4*<sup>-/-</sup> + EtOH | p = 0.0093 |  |  |  |  |

**Supplemental Figure 1.**
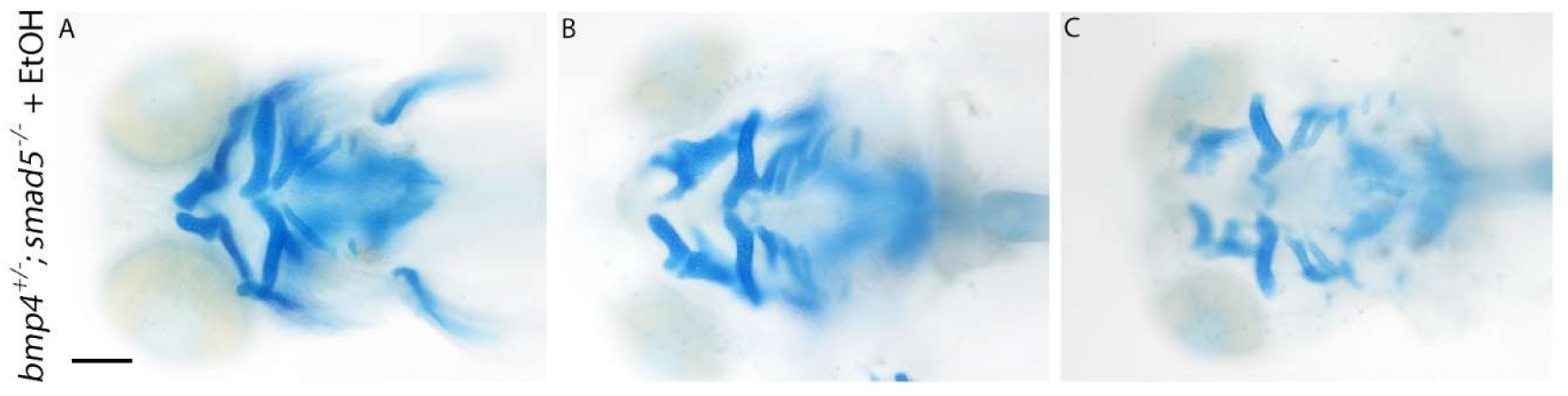
A wide spectrum of viscerocranial malformations was observed in ethanol-treated *bmp4^+/-^;smad5^-/-^* larvae. (**A-C**) Whole-mount images of viscerocranium at 5 dpf larvae. Cartilage is blue and bone is red (Ventral views, anterior to the left, scale bar: 100 μm). (**A-C**) Ethanol exposure on *bmp4^+/-^;smad5^-/-^*larvae range from stereotypical *smad5* mutant phenotypes to severe ethanol-induced phenotypes to the viscerocranium.

**Supplemental Table 4.**
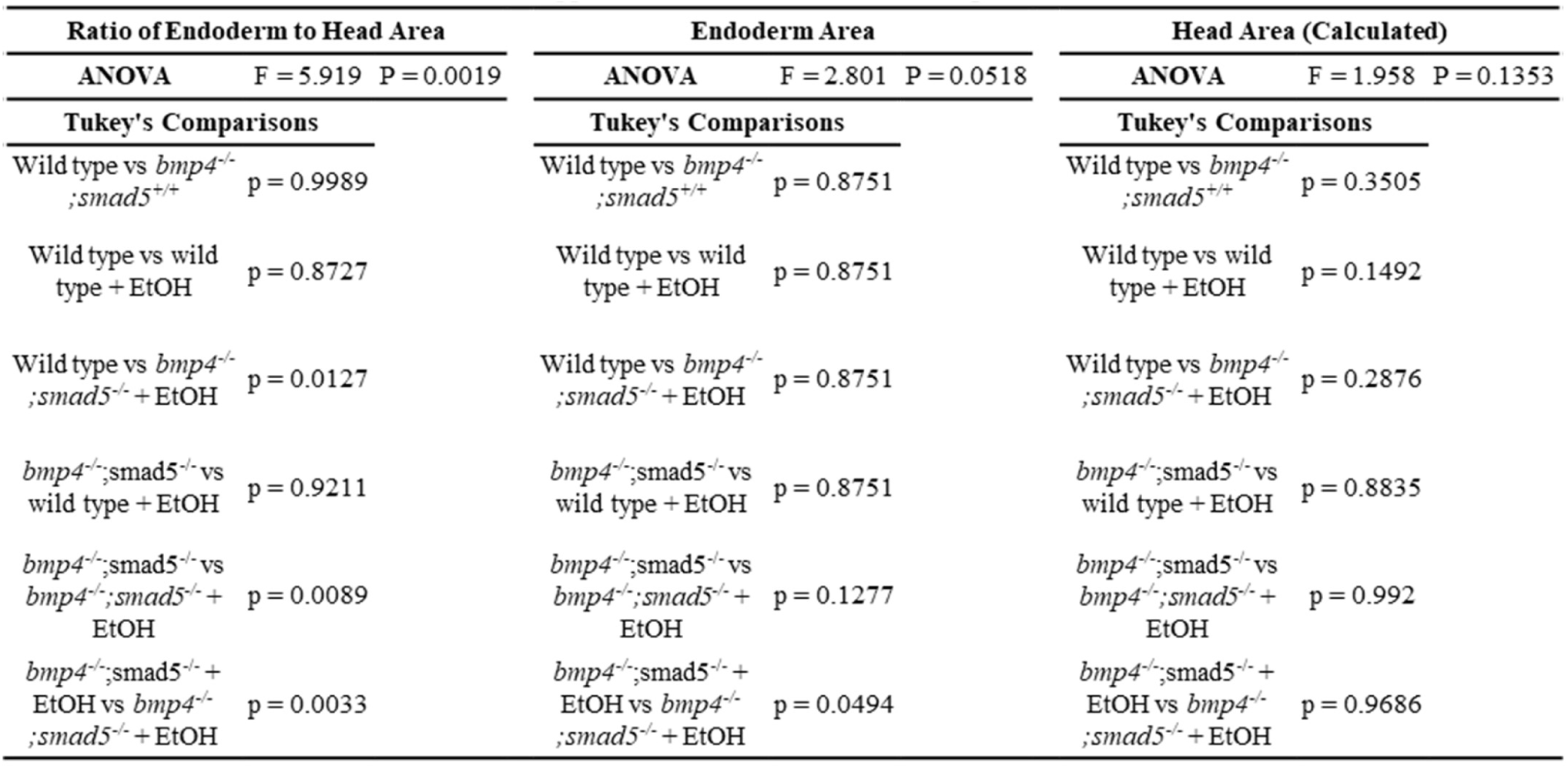
Statistical analyses of endoderm/head area measures in Figure 4. Measures of anterior endoderm/head area were analyzed with a two-way ANOVA (type III). F-statistic and P-value for each analysis are shown. A Tukey’s Multiple Comparisons Test for each comparison are shown.

**Supplemental Figure 2.**
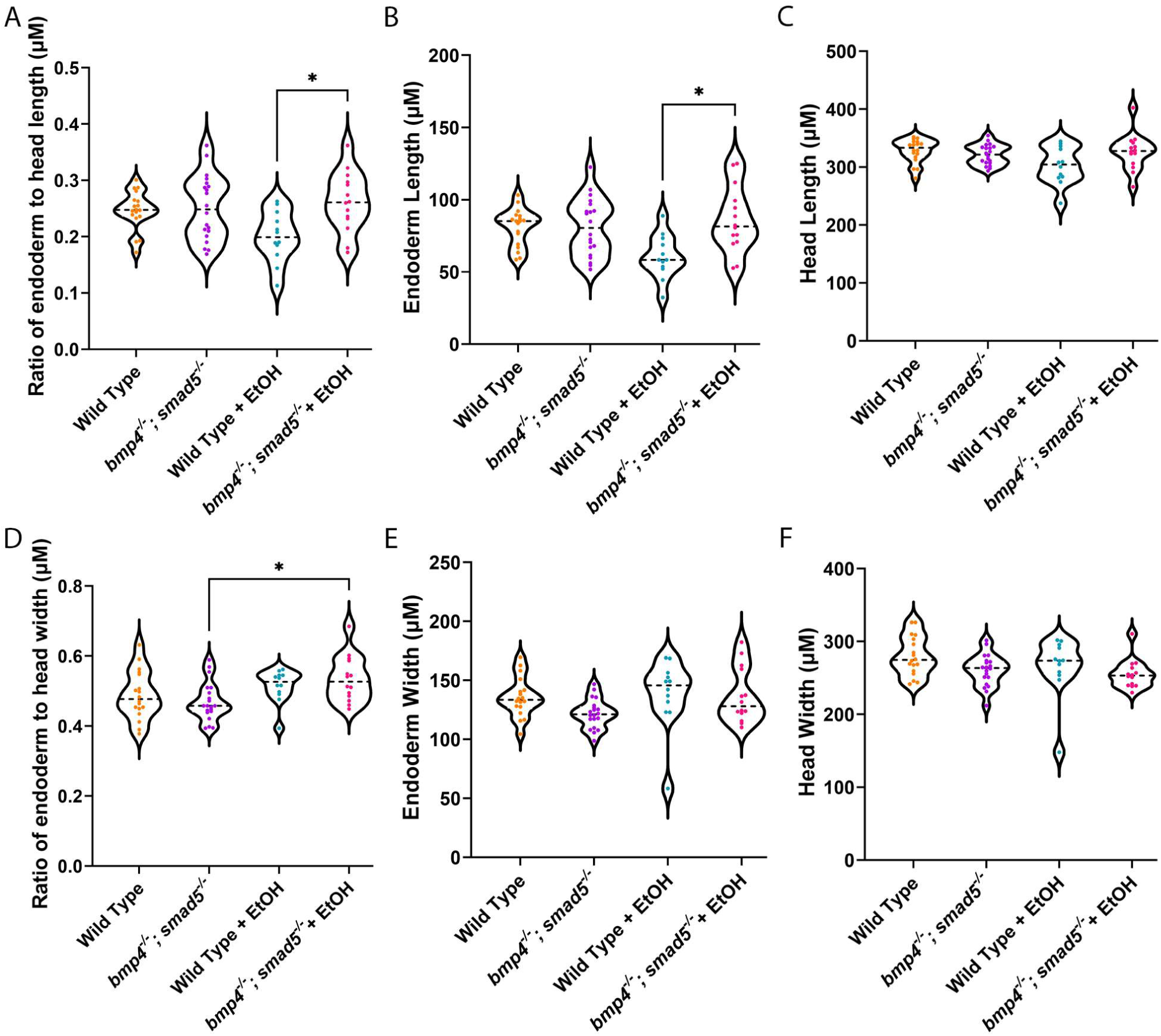
Ethanol exposure alters size of the anterior pharyngeal endoderm in *bmp4^-/-^;smad5^-/-^* embryos. (**A-F**) Length and width measures of the anterior pharyngeal endoderm. (**A-C**) Ethanol exposure decreased endoderm length but not head length in Wild Type embryos but significantly increased endoderm length ethanol-treated *bmp4^-/-^;smad5^-/-^* embryos. (**D-F**) Ethanol-treatment significantly increased in the ratio of endoderm width to head width in *bmp4^-/-^;smad5^-/-^* embryos, though there was a trend in the increase in ethanol-treated Wild Type embryos as well, but not in head width (individual graph statistics in Supplemental Table 3).

**Supplemental Table 5.** Statistical analyses of linear anterior endoderm/head measures in Supplemental Figure 2. Linear measures of anterior endoderm/head were analyzed with a two-way ANOVA (type III). F-statistic and P-value for each analysis are shown. A Tukey’s Multiple Comparisons Test for each comparison are shown.

| Ratio of Endoderm to Head Length |  | Endoderm Length |  | Head Length |  |
| --- | --- | --- | --- | --- | --- |
| ANOVA | F = 2.930 P = 0.0448 | ANOVA | F = 3.811 P = 0.0169 | ANOVA | F = 2.309 P = 0.0906 |
| Tukey's Comparisons |  | Tukey's Comparisons |  | Tukey's Comparisons |  |
| Wild type vs *bmp4*<sup>-/-</sup>;<br>*smad5*<sup>+/+</sup> | p = 0.9989 | Wild type vs *bmp4*<sup>-/-</sup>;<br>*smad5*<sup>+/+</sup> | p = 0.8751 | Wild type vs *bmp4*<sup>-/-</sup>;<br>*smad5*<sup>+/+</sup> | p = 0.3505 |
| Wild type vs wild<br>type + EtOH | p = 0.8727 | Wild type vs wild<br>type + EtOH | p = 0.8751 | Wild type vs wild<br>type + EtOH | p = 0.1492 |
| Wild type vs *bmp4*<sup>-/-</sup>;<br>*smad5*<sup>-/-</sup> + EtOH | p = 0.0127 | Wild type vs *bmp4*<sup>-/-</sup>;<br>*smad5*<sup>-/-</sup> + EtOH | p = 0.8751 | Wild type vs *bmp4*<sup>-/-</sup>;<br>*smad5*<sup>-/-</sup> + EtOH | p = 0.2876 |
| *bmp4*<sup>-/-</sup>;*smad5*<sup>-/-</sup> vs<br>wild type + EtOH | p = 0.9211 | *bmp4*<sup>-/-</sup>;*smad5*<sup>-/-</sup> vs<br>wild type + EtOH | p = 0.8751 | *bmp4*<sup>-/-</sup>;*smad5*<sup>-/-</sup> vs<br>wild type + EtOH | p = 0.8835 |
| *bmp4*<sup>-/-</sup>;*smad5*<sup>-/-</sup> vs<br>*bmp4*<sup>-/-</sup>;*smad5*<sup>-/-</sup> +<br>EtOH | p = 0.0089 | *bmp4*<sup>-/-</sup>;*smad5*<sup>-/-</sup> vs<br>*bmp4*<sup>-/-</sup>;*smad5*<sup>-/-</sup> +<br>EtOH | p = 0.1277 | *bmp4*<sup>-/-</sup>;*smad5*<sup>-/-</sup> vs<br>*bmp4*<sup>-/-</sup>;*smad5*<sup>-/-</sup> +<br>EtOH | p = 0.992 |
| *bmp4*<sup>-/-</sup>;*smad5*<sup>-/-</sup> +<br>EtOH vs *bmp4*<sup>-/-</sup>;<br>*smad5*<sup>-/-</sup> + EtOH | p = 0.0033 | *bmp4*<sup>-/-</sup>;*smad5*<sup>-/-</sup> +<br>EtOH vs *bmp4*<sup>-/-</sup>;<br>*smad5*<sup>-/-</sup> + EtOH | p = 0.0494 | *bmp4*<sup>-/-</sup>;*smad5*<sup>-/-</sup> +<br>EtOH vs *bmp4*<sup>-/-</sup>;<br>*smad5*<sup>-/-</sup> + EtOH | p = 0.9686 |
| Ratio of Endoderm to Head Width |  | Endoderm Width |  | Head Width |  |
| ANOVA | F = 3.063 P = 0.0386 | ANOVA | F = 1.869 P = 0.1499 | ANOVA | F = 2.691 P = 0.0587 |
| Tukey's Comparisons |  | Tukey's Comparisons |  | Tukey's Comparisons |  |
| Wild type vs *bmp4*<sup>-/-</sup>;<br>*smad5*<sup>+/+</sup> | p = 0.8133 | Wild type vs *bmp4*<sup>-/-</sup>;<br>*smad5*<sup>+/+</sup> | p = 0.2316 | Wild type vs *bmp4*<sup>-/-</sup>;<br>*smad5*<sup>+/+</sup> | p = 0.1886 |
| Wild type vs wild<br>type + EtOH | p = 0.652 | Wild type vs wild<br>type + EtOH | p = 0.9983 | Wild type vs wild<br>type + EtOH | p = 0.3263 |
| Wild type vs *bmp4*<sup>-/-</sup>;<br>*smad5*<sup>-/-</sup> + EtOH | p = 0.1846 | Wild type vs *bmp4*<sup>-/-</sup>;<br>*smad5*<sup>-/-</sup> + EtOH | p = 0.9993 | Wild type vs *bmp4*<sup>-/-</sup>;<br>*smad5*<sup>-/-</sup> + EtOH | p = 0.0523 |
| *bmp4*<sup>-/-</sup>;*smad5*<sup>-/-</sup> vs<br>wild type + EtOH | p = 0.2343 | *bmp4*<sup>-/-</sup>;*smad5*<sup>-/-</sup> vs<br>wild type + EtOH | p = 0.277 | *bmp4*<sup>-/-</sup>;*smad5*<sup>-/-</sup> vs<br>wild type + EtOH | p > 0.9999 |
| *bmp4*<sup>-/-</sup>;*smad5*<sup>-/-</sup> vs<br>*bmp4*<sup>-/-</sup>;*smad5*<sup>-/-</sup> +<br>EtOH | p = 0.0324 | *bmp4*<sup>-/-</sup>;*smad5*<sup>-/-</sup> vs<br>*bmp4*<sup>-/-</sup>;*smad5*<sup>-/-</sup> +<br>EtOH | p = 0.2553 | *bmp4*<sup>-/-</sup>;*smad5*<sup>-/-</sup> vs<br>*bmp4*<sup>-/-</sup>;*smad5*<sup>-/-</sup> +<br>EtOH | p = 0.8529 |
| *bmp4*<sup>-/-</sup>;*smad5*<sup>-/-</sup> +<br>EtOH vs *bmp4*<sup>-/-</sup>;<br>*smad5*<sup>-/-</sup> + EtOH | p = 0.8716 | *bmp4*<sup>-/-</sup>;*smad5*<sup>-/-</sup> +<br>EtOH vs *bmp4*<sup>-/-</sup>;<br>*smad5*<sup>-/-</sup> + EtOH | p > 0.9999 | *bmp4*<sup>-/-</sup>;*smad5*<sup>-/-</sup> +<br>EtOH vs *bmp4*<sup>-/-</sup>;<br>*smad5*<sup>-/-</sup> + EtOH | p = 0.866 |

**Supplemental Figure 3.**
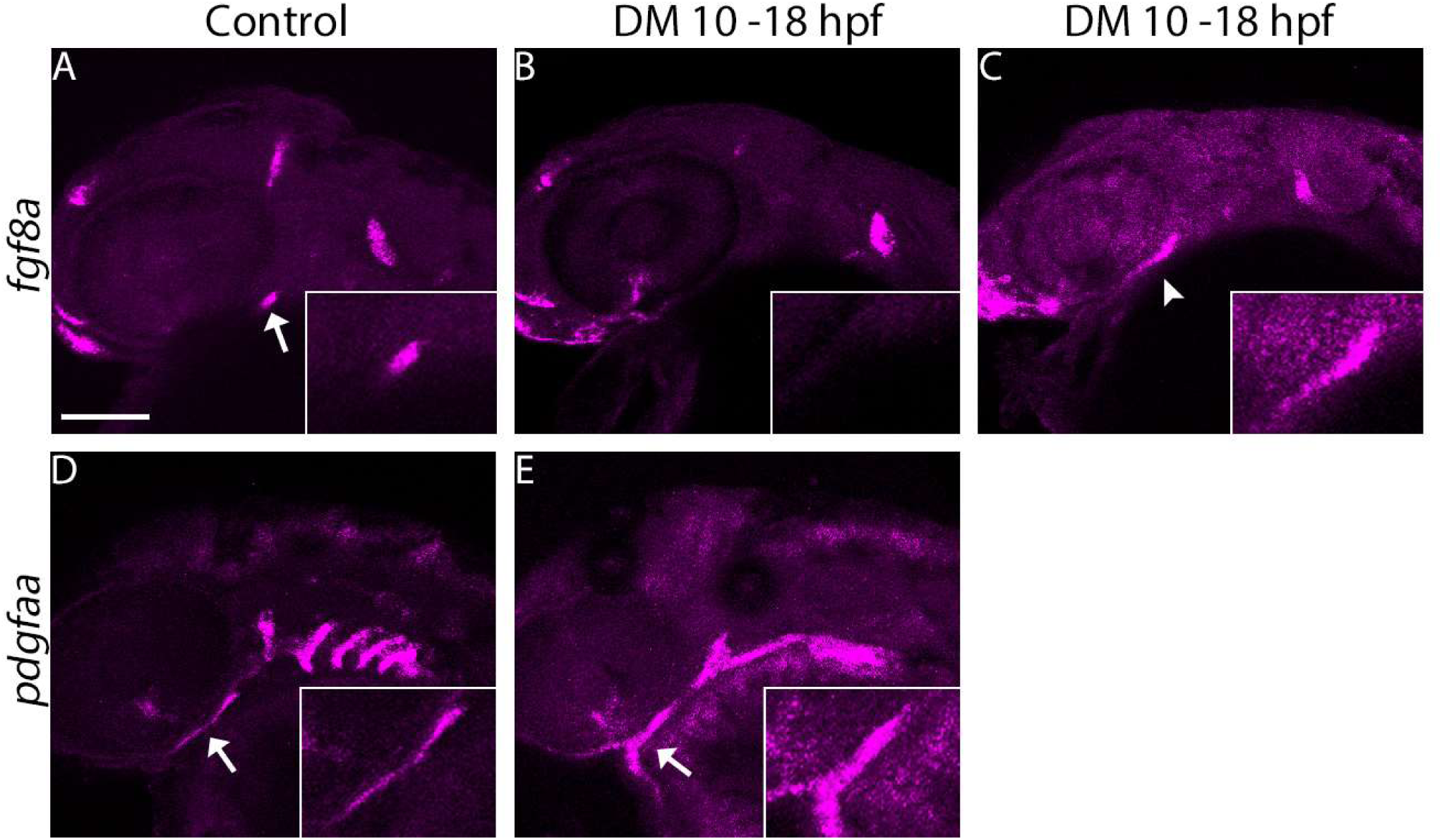
Blocking Bmp signaling with Dorsomorphin inhibitor (DM) disrupts the expression of *fgf8a* in the oral ectoderm. (**A-F**) Whole-mount images of untreated and DM-treated embryos labeling *fgf8a* and *pdgfaa* gene expression at 36 hpf (lateral views, anterior to the left, scale bar: 100 μm). (**A&D**) Normal expression of *fgf8a* and *pdgfaa* in the oral ectoderm of untreated embryos. (**B, C, E**) Expression of *fgf8a* is lost in DM-treated embryos, while *pdgfaa* are expressed normally (n = 20 embryos per group).

**Supplemental Figure 4.**
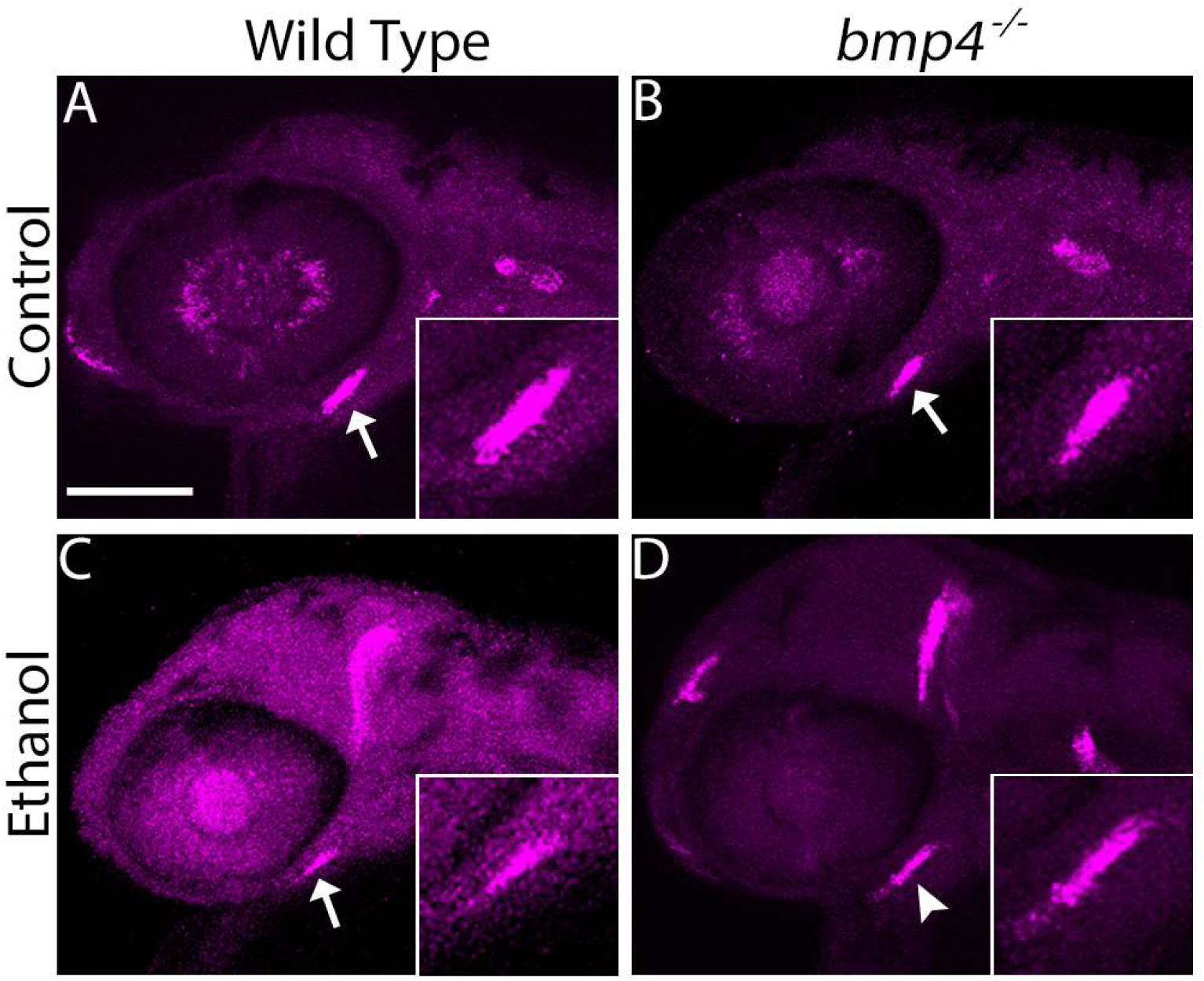
Ethanol exposure changes shape of oral ectoderm expression domain in *bmp4^-/-^* embryos. (**A-D**) Whole-mount, confocal images of *bmp4* embryos fluorescently labeling *fgf8a* gene expression at 36 hpf (lateral views, anterior to the left, scale bar: 100 μm). (**A-C**) Arrows show normal expression of *fgf8a* in the oral ectoderm of untreated wild type and *bmp4^-/-^* embryos as well as ethanol-treated wild type embryos. (**D**) Arrowhead shows that domain of *fgf8a* expression in ethanol-treated *bmp4^-/-^* embryos is subtly expanded anteriorly (n = 5 embryos per group).

**Supplemental Figure 5.**
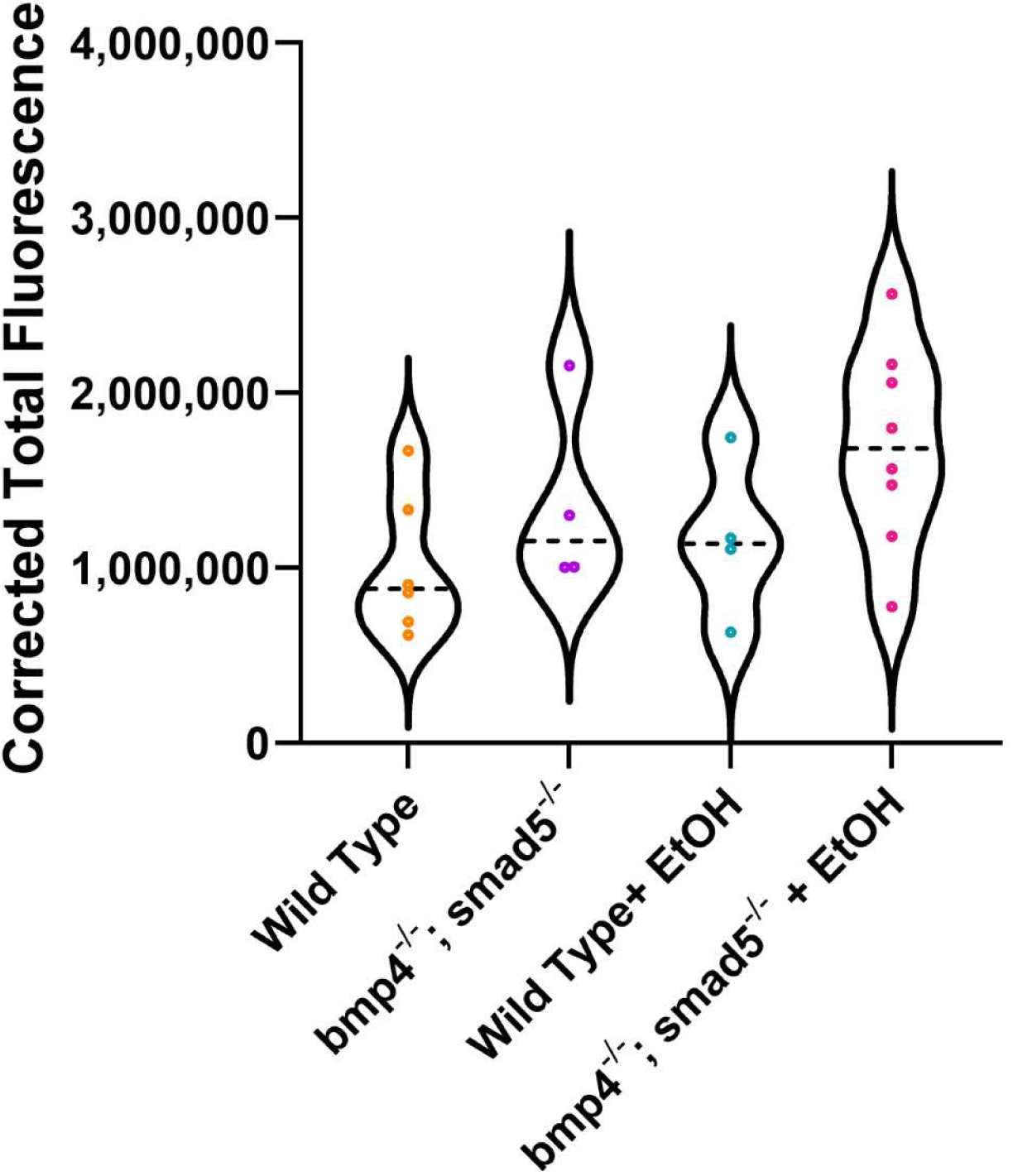
Ethanol exposure does not lead to decreases in Bmp signaling responses. Corrected Total Fluorescence was calculated from *BRE:mKO2* fluorescence. We observed no change in BRE signaling responses in *bmp4^-/-^;smad5^-/-^* embryos or due to ethanol. (Embryos per group, wild type, n = 5; *bmp4^-/-^;smad5^-/-^*, n = 4; wild type + Etoh, n = 4; *bmp4^-/-^;smad5^-/-^*, n = 8).

**Supplemental Table 6.**
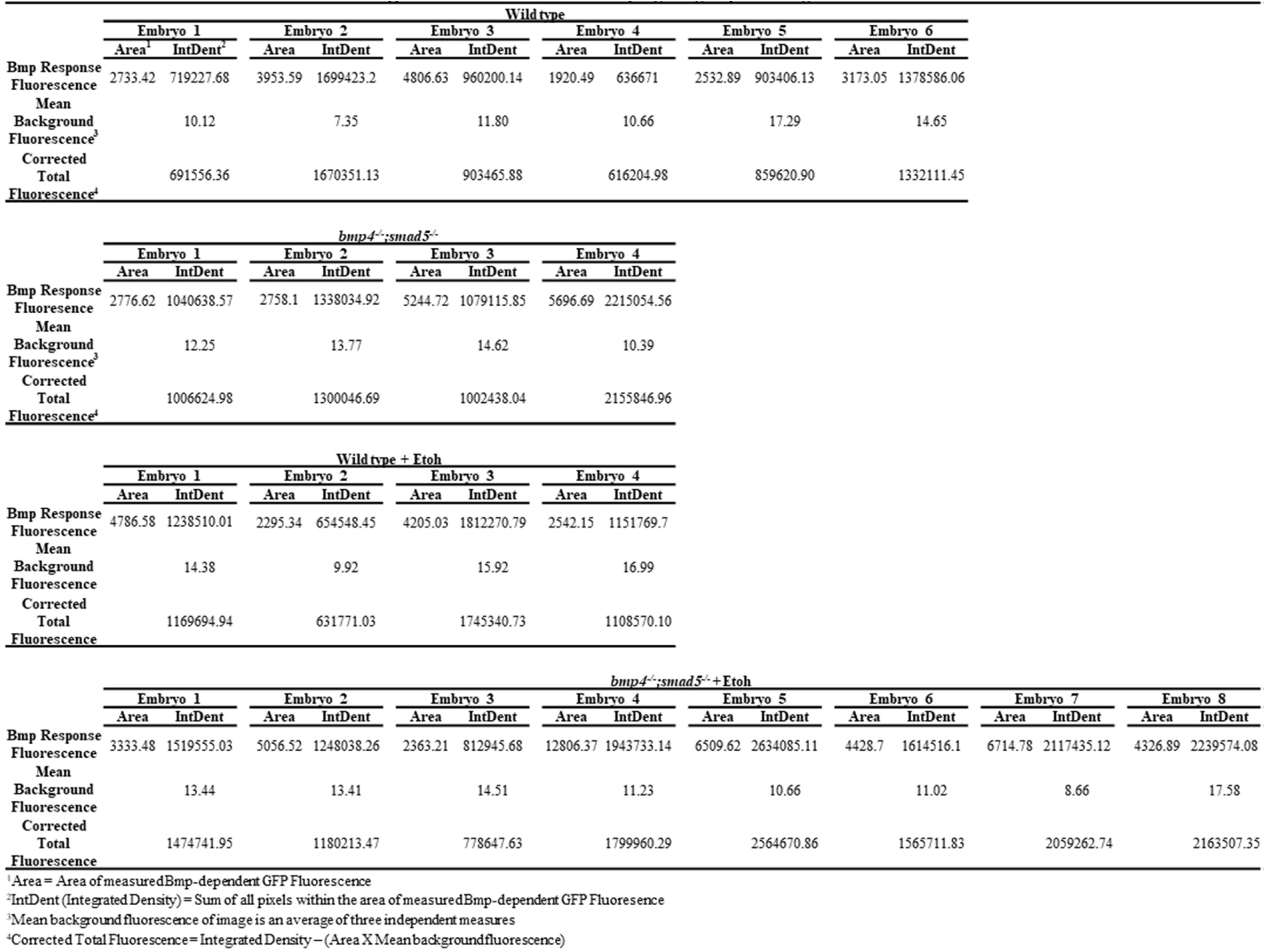
Calculation of Corrected Total Fluorescence for Figures 6 & S5. Pharyngeal area of *BRE:mKO2* fluorescent intensity was quantified using Image J. Corrected Total Fluorescence (unitless) was calculated from Integrated Density – (Area of fluorescence x mean background fluorescence). Integrated density is the Sum of all pixels within the area of fluorescent measurement. Mean background fluorescence was the average of three independent measures.

**Supplemental Figure 6.**
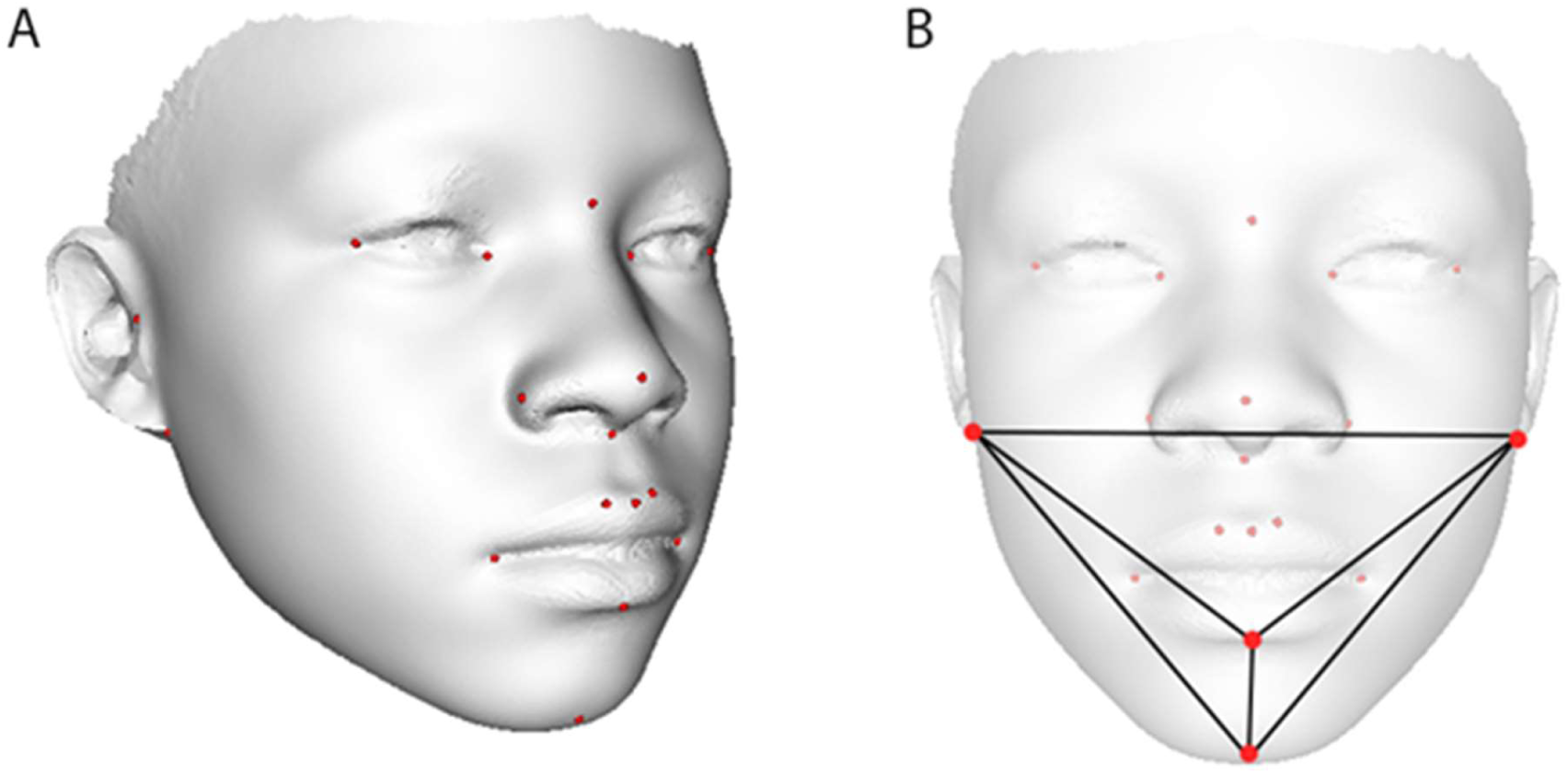
Visual representation of the 20 landmark locations used to induce the dense surface correspondence. (**A-B**) left and right endocanthion, exocanthion, tragion, otobasion inferius, crista philtrum, cheilion, and alare; nasion, pronasale, subnasale, labiale superius, labiale inferius, and gnathion. (**B**) The landmarks labiuminferius, gnathion, and otobasion inferior define the tetrahedron (1,2,3,4). The volume of the tetrahedron is calculated as (1/6)det (axayaz1; bxbybz1; cxcycz1; dxdydz1).

